# Isolation of a natural DNA virus of *Drosophila melanogaster*, and characterisation of host resistance and immune responses

**DOI:** 10.1101/215111

**Authors:** William H. Palmer, Nathan Medd, Philippa M. Beard, Darren J. Obbard

## Abstract

*Drosophila melanogaster* has played a key role in our understanding of invertebrate immunity. However, both functional and evolutionary studies of host-virus interaction in *Drosophila* have been limited by a dearth of native virus isolates. In particular, despite a long history of virus research, DNA viruses of *D. melanogaster* have only recently been described, and none have been available for experimental study. Here we report the isolation and comprehensive characterisation of Kallithea virus, a large double-stranded DNA virus, and the first DNA virus to have been reported from wild populations of *D. melanogaster*. We find that Kallithea virus infection is costly for adult flies, reaching high titres in both sexes and disproportionately reducing survival in males and movement and late fecundity in females. Using the *Drosophila* Genetic Reference Panel, we quantify host genetic variance for virus-induced mortality and viral titre and identify candidate host genes that may underlie this variation, including *Cdc42-interacting protein 4*. Using full transcriptome sequencing of infected males and females, we examine the transcriptional response of flies to Kallithea virus infection, and describe differential regulation of virus-responsive genes. This work establishes Kallithea virus as a new tractable model to study the natural interaction between *D. melanogaster* and DNA viruses, and we hope it will serve as a basis for future studies of immune responses to DNA viruses in insects.

**Author Summary:** The fruit fly *Drosophila melanogaster* is a useful model species to study host-virus interaction and innate immunity. However, few natural viruses of *Drosophila* have been available for experiments, and no natural DNA viruses of *Drosophila melanogaster* have been available at all. Although infecting flies with viruses from other insects has been useful to uncover general immune mechanisms, viruses that naturally infect wild flies could help us to learn more about the coevolutionary process, and more about the genes that underlie the host-virus interaction. Here we present an isolate of a DNA virus (named Kallithea Virus) that naturally infects the model species *Drosophila melanogaster* in the wild. We describe the basic biology of infection by this virus, finding that both male and females flies die from infection, but females are more tolerant of infection than males, while laying lay fewer eggs than uninfected females. We quantify genetic variation for virus resistance in the flies, and we use RNA sequencing to see which genes are expressed in male and female flies in response to infection. These results will form the basis for further research to understand how insects defend themselves against infection by DNA viruses, and how DNA viruses can overcome antiviral defence.

## Introduction

Studies of *Drosophila melanogaster* are central to our understanding of infection and immunity in insects. Moreover, many components of the *Drosophila* immune response, including parts of the JAK-STAT, IMD, and Toll (and perhaps RNA interference; RNAi) pathways are conserved from flies to mammals (Dupuis et al, 2003; Karst et al, 2003; Sharma et al, 2003; Zambon et al, 2005; Dostert et al, 2005; Wang et al, 2006; Avadhanula et al, 2009; Maillard et al, 2013; Li et al, 2013), making *Drosophila* a valuable model beyond the insects. The experimental dissection of antiviral immune pathways in *Drosophila* has benefited from both natural infectious agents of *Drosophila*, such as Drosophila C Virus (DCV) and Sigma virus (DmelSV), and from artificial infections, such as Cricket paralysis virus (isolated from a cricket), Flock House Virus (from a beetle), Sindbis virus (from a mosquito) and Invertebrate Iridescent Virus 6 (from a moth). However, while the availability of experimentally tractable, but non-natural, model viruses has been a boon to studies of infection, it also has two potential disadvantages. First, the coevolutionary process means that pairs of hosts and pathogens that share a history may interact very differently to naive pairs (e.g. Ferguson and Read, 2002; Compton et al, 2012). For example, the Nora virus of *D. immigrans* expresses a viral suppressor of RNAi that is functional in the natural host, but not in *D. melanogaster* (van Mierlo et al, 2014). Second, if our aim is to understand the coevolutionary process itself, then the standing diversity in both host and virus populations may be fundamentally altered in coevolving as opposed to naïve pairs. For example, heritable variation for host resistance was detectable for two natural viruses of *D. melanogaster*, but not for two non-natural viruses (Magwire et al, 2012; Wang et al, 2017). This difference was in part due to large-effect segregating polymorphisms for resistance to the natural viruses, which are predicted to result from active coevolutionary dynamics (Contamine et al, 1989; Magwire et al, 2011; Magwire et al, 2012; Cogni et al, 2016).

Experimental studies of host-virus interaction using *Drosophila* have consequently been limited by a lack of diverse natural virus isolates. In particular, no natural DNA viral pathogens of *D. melanogaster* have previously been isolated (Brun and Plus, 1980; Huszar and Imler, 2008; but see Unckless, 2011 for a DNA virus of *Drosophila innubila*), and all natural (and most artificial) studies of viral infection in *D. melanogaster* have therefore focussed on the biology of RNA viruses and antiviral resistance (Xu and Cherry, 2014; Bronkhorst et al, 2012). For DNA viruses, our molecular understanding of insect-virus interaction has instead largely been shaped by the response of lepidopterans to their natural Baculoviruses, which are often of agronomic and/or ecological importance (Herniou et al, 2004), but lack the genetic toolkit of *D. melanogaster*. Nevertheless, Lepidopteran expression studies of the expression response to baculovirus infection have implicated host genes with a diverse array of functions, including cuticle proteins, reverse transcriptases, and apoptotic factors, suggesting previously uncharacterised and/or host-specific antiviral immune mechanisms (Breitenbach et al, 2011; Noland et al, 2013; Nguyen et al, 2013; McTaggart et al 2015).

To date, the only DNA virus studies in *D. melanogaster* have used Insect Iridescent Virus 6 (IIV6), an enveloped dsDNA moth iridovirus with an extremely broad host range (Williams 2008). This work has shown that *Drosophila* RNAi mutants are hyper-susceptible to IIV6 infection, and that IIV6 encodes a viral suppressor of RNAi, indicating that at least some immune responses to DNA viruses overlap substantially with those to RNA viruses (Kemp et al, 2013; Bronkhorst et al, 2012; Bronkhorst et al, 2014). However, while IIV6 injections are lethal in *D. melanogaster*, and IIV6 has provided useful information about the *Drosophila* response to DNA viruses, for the reasons described above, it is hard to interpret the implications of this for our understanding of natural host-virus interaction.

Metagenomic sequencing has identified natural dsDNA nudivirus infections in wild-caught *D. innubila* (*D. innubila* Nudivirus, DiNV; Unckless, 2011; Hill and Unckless, 2017) and in *D. melanogaster* and *D. simulans* (‘Kallithea virus’, KV; Webster et al, 2015; ‘Esparto virus’ (KY608910.1), and ‘Tomelloso virus’ (KY457233.1)), and also ssDNA densovirus infections in *D. melanogaster* and *D. simulans* (‘Vesanto virus’ (KX648534.1), ‘Linvill Road virus’ (KX648536.1), and ‘Viltain virus’ (KX648535.1)). Like other members of the *Nudiviridae*, DiNV and KV are enveloped dsDNA viruses of around 120-230Kbp with 100-150 genes. This recently-recognised family forms a clade that is either sister to, or paraphyletic with, the Bracoviruses (Thézé et al, 2011) that have been ‘domesticated’ by Braconid parasitoid wasps following genomic integration, and now provide key components of the wasp venom (Herniou et al, 2013; Gauthier et al, 2017). Together, the nudiviruses and bracoviruses are sister to the baculoviruses, which are arguably the best-studied dsDNA viruses of insects. They share many of their core genes with baculoviruses, but canonically lack occlusion bodies (Wang and Jehle, 2009). PCR surveys of wild flies suggest that DiNV is common in several species in the subgenus *Drosophila*, and that KV is widespread and common in *D. melanogaster* and *D. simulans*, being detectable in 10 of 17 tested populations, with an estimated global prevalence of 2-7% (Webster et al, 2015). However, we currently know little about the interaction between these viruses and their hosts. Indeed, although studies of wild-caught *D. innubila* individuals infected by DiNV suggest that infection is costly (Unckless 2011), in the absence of an experimental *D. melanogaster* nudivirus isolate, it has not been possible to capitalise the power of *D. melanogaster* genetics to further elucidate the costs associated with infection, or the genetic basis of resistance.

Here we present the isolation of KV from wild-collected *D. melanogaster* via passage in laboratory stocks and gradient centrifugation. We use this isolate to characterise the fundamental phenotypic impacts of infection on host longevity and fecundity. We then use the *Drosophila* Genetic Reference Panel (DGRP) to quantify and dissect genetic variation in immunity to KV infection in males and females, and we use RNAseq analyses of an inbred line to quantify host and virus transcriptional response in both sexes. We find that KV causes higher rates of mortality following injection in males, but that male and female viral titres do not differ substantially, suggesting some female tolerance to infection. However, we also find that female movement is decreased following infection, and that infected females have significantly reduced late-life fecundity—highlighting the importance of considering infection phenotypes beyond longevity. We find a genetic correlation in longevity between KV-infected males and females, and a weak negative genetic correlation between mortality and KV titre in females, and we report host loci that have variants significantly associated with each trait. Finally, our expression analysis of infected individuals supports a dramatic cessation of oogenesis following infection, and significant differential regulation of serine proteases and certain immune genes. This work establishes KV as a new natural model for DNA virus infection in *D. melanogaster* and will enable further dissection of the insect antiviral immune response.

## Materials and Methods

### Isolation of Kallithea Virus

We identified KV-infected flies through a PCR screen for previously published *D. melanogaster* viruses in 80 previously untested wild-caught flies (see Webster et al, 2015 for primers and cycling conditions). We homogenised each fly in 0.1 mL of Ringer’s solution, transferred half of the homogenate to Trizol for nucleic acid extraction, and performed RT PCR assays on the resulting RNA for all *D. melanogaster* viruses reported by Webster *et al* (2015). We selected a KV-positive sample from Thika, Kenya (Collected by John Pool in 2009; subsequently stored at -80C), removed debris from the remaining fly homogenate by centrifugation for 10 minutes at 1000 × g, and microinjected 50 nL of the supernatant into *Dicer-2^L811fsX^* flies, which lack a robust antiviral immune response (Lee et al, 2004). After one week, we homogenised 100 KV-injected *Dicer-2 ^L811fsX^* flies in 10 uL Ringer’s solution per fly, cleared the solution by centrifugation as above, and re-injected this homogenate into further *Dicer-2^L811fsX^* flies. This process was then repeated twice more with the aim of increasing viral titres. In the final round of serial passage, we injected 2000 *Dicer-2 ^L811fsX^* flies, which were homogenised in 5 mL 10 mM Tris-HCl. We cleared the homogenate by centrifuging at 1000 × g for 10 minutes, filtering through cheese cloth, centrifuging twice more at 6000 × g for 10 minutes, and finally filtering through a Millex 0.45 µm polyvinylidene fluoride syringe filter. The resulting crude virus preparation was used as input for gradient ultracentrifugation.

We screened the crude preparation by RT-PCR for other published *Drosophila* virus sequences, and identified the presence of DAV, Nora virus, DCV, and La Jolla virus. To separate KV from these viruses, we used equilibrium buoyant density centrifugation in iodixanol (“OptiPrep”, Sigma-Aldrich) as enveloped viruses are expected to have lower buoyant densities than most unenveloped viruses. Iodixanol is biologically inert, and gradient fractions can be used directly for downstream infection experiments (avoiding dialysis, which we found greatly reduces KV titres). We concentrated virus particles by centrifuging crude virus solution through a 1 mL 10% iodixanol layer onto a 2 mL 30% iodixanol cushion at 230,000 × g for 4 hours in a Beckman SW40 rotor. Virus particles were taken from the 30%-10% interphase, and layered onto a 40%-10% iodixanol step gradient, with 2% step changes, and centrifuged for 48 hours at 160,000 × g. We fractionated the gradient at 0.5 mL intervals, phenol-chloroform extracted total nucleic acid from aliquots of each fraction, and measured virus concentration by quantitative PCR (qPCR). We pooled all Kallithea-positive, RNA virus-negative fractions and calculated the infectious dose 50 (ID50) by injecting 3 vials of 10 flies with a series of 10-fold dilutions and performing qPCR after 5 days. We simultaneously performed the above isolation protocol with uninfected *Dicer-2^L811fsX^* flies, and extracted the equivalent fractions for use as an injection control solution (hereafter referred to as “gradient control”).

### Transmission electron microscopy

A droplet of viral suspension was allowed to settle on a Formvar/Carbon 200 mesh Copper grid for 10 minutes. We removed excess solution and applied a drop of 1% aqueous uranyl acetate for 1 minute before removing the excess by touching the grid edge with filter paper. The grids were then air dried. Samples were viewed using a JEOL JEM-1400 Plus transmission electron microscope, and representative images were collected on a GATAN OneView camera.

### Measurement and analysis of viral titre

Flies were reared on a standard cornmeal diet until infection, after which they were transferred to a solid sucrose-agar medium. We infected flies by abdominal injection of 50 nL of 10^5^ ID50 KV using a Nanoject II (Drummond Scientific), and these flies were then used to assay changes in viral titre, mortality, fecundity, or daily movement. To test whether viral titre over time was influenced by sex or the presence of *Wolbachia* endosymbionts, we injected 25 vials of 10 male or female *Oregon R* flies with KV, with or without *Wolbachia* (totalling 1000 flies). We phenol-chloroform extracted total nucleic acid at 5 time points: directly after injection and 3, 5, 10, and 15 days post-infection. We used qPCR to measure viral titre relative to copies of the fly genome with the following (PCR primers: kallithea_126072F CATCAATATCGCGCCATGCC, kallithea_126177R GACCGAGTTAGCGTCAATGC, rpl32_465F CTAAGCTGTCGGTGAGTGCC, rpl32_571R: TGTTGTCGATACCCTTGGGC). We analysed the log-transformed relative expression levels of Kallithea virus as a Gaussian response variable in a linear mixed model using the Bayesian generalised mixed modelling R package MCMCglmm (V2.24; Hadfield, 2010, see supporting text for code to fit all models).

The fixed effects portion of the model included an intercept term and coefficients for the number of days post-inoculation (DPI), sex, and DPI by sex interaction. We estimated random effects for each qPCR plate, and assumed random effects and residuals are normally distributed. We initially fitted the model with *Wolbachia* infection status included as a fixed effect, however this term was not significant and was excluded from the final model.

We also attempted to infect flies with KV by feeding. We anesthetised flies in an agar vial and sprayed 50 uL of 5×10^3^ ID50 KV onto the flies and food. We then collected flies immediately (for the zero time-point) and at 7 DPI and used the primers above to calculate relative KV titre.

### Mortality following KV infection

We performed mortality assays to test the effect of KV infection on longevity, and to test whether this was affected by sex or *Wolbachia* infection status. We injected a total of 1200 *Oregon R* flies with control gradient or KV for each sex with or without *Wolbachia* (*Wolbachia* had previously been cleared by 3 generations of Ampicillin treatment and its absence was confirmed by PCR). We maintained flies for each treatment in 10 vials of 10 flies, and recorded mortality daily for three weeks. Mortality that occurred in the first day after infection was assumed to be due to the injection procedure and excluded from further analysis. We analysed mortality using an event-analysis framework as a generalised linear mixed model in MCMCglmm, with per-day mortality in each vial as a binomial response variable. We included fixed effects for DPI, DPI^2^ (used to capture nonlinear mortality curves), KV infection status, the two-way interaction between DPI and KV infection status, the two-way interaction between DPI and sex, and the three-way interaction between DPI, KV infection status, and sex. We fitted vials as a random effect to account for non-independence among flies within vials, assuming these follow a normal distribution. As in the model for viral titre, we found no evidence for differences associated with *Wolbachia* infection, and *Wolbachia* terms were excluded from the final model. The higher rate of male mortality we observed was also confirmed in a second independent experiment using an outbred population derived from the Drosophila Genetic Reference Panel (DGRP; see below).

### Fecundity following KV infection

We measured fecundity during early (1 and 2 DPI) and late (7 and 8 DPI) Kallithea virus infection. Virgin female flies from an outbred population derived from the DGRP (Mackay et al, 2012; created from 113 DGRP lines and maintained at a low larval density with non-overlapping generations) were injected with either KV, or with chloroform-inactivated KV as a control, and individually transferred to standard cornmeal vials. The following day we introduced a single male fly into the vial with the virgin female. We transferred the pair to new vials each day and recorded the number of eggs laid. Per-day fecundity was analysed in MCMCglmm as a Poisson response variable using a hurdle model, which models the probability of zeroes in the data and the Poisson process as separate variables. We included fixed effects associated with KV infection status, infection stage (early or late), the interaction between KV infection and infection stage, and random effects associated with each fly pair (vial).

We analysed ovary morphology to examine whether changes in fecundity were detectable in ovaries. Flies were injected with either control virus solution or KV and kept on agar vials. After 8 DPI, flies were transferred to vials with Lewis medium supplemented with yeast. Two days later, we dissected ovaries in phosphate-buffered saline solution, fixed ovaries in 4% paraformaldehyde, and stained nuclei with DAPI. Ovaries were analysed under a Leica fluorescence microscope, and we recorded whether each ovariole within an ovary included egg chambers past stage 8 (i.e. had begun vitellogenesis), and whether any egg chambers within an ovariole exhibited apoptotic nurse cells. The probability of an ovariole containing a post-vitellogenic egg chamber was analysed using a logistic regression in MCMCglmm, with KV infection status as a fixed effect and the ovary from which the ovariole derived as a random effect. We analysed whether apoptotic nurse cells are associated with KV virus-infected ovary in a similar manner.

### Daily movement following KV infection

We used a *Drosophila* Activity Monitor (DAM, TriKinetics; Pfeiffenberger et al, 2010) to measure per-day total movement of individual flies (Pfeiffenberger et al, 2010). The DAM is composed of multiple hubs, each with 32 tubes containing a single fly, and movement is recorded on each occasion the fly breaks a light beam. We injected 96 female flies from an outbred DGRP population with either chloroform-inactivated KV or KV, randomly assigned these flies within and across 3 hubs, and measured total movement for one week. Movement was binned for each day and this per-day total movement was analysed in a linear mixed model using MCMCglmm as a Poisson response variable. We completely excluded flies that failed to move for a whole day or longer, assuming them to be dead. As before, we included fixed effects associated with DPI, KV infection status, and the interaction between KV and DPI. We included random effects associated with each fly (repeated measures) and each of the DAM hubs, and assumed each of these take values from a normal distribution.

### Quantitative genetic analysis

The DGRP is a collection of highly inbred fly lines derived from a *D. melanogaster* population collected in Raleigh, North Carolina (Mackay et al, 2012), and is widely used to estimate and dissect genetic variation in complex traits in *Drosophila*. We measured KV titre in females and mortality following KV infection in both sexes for 125 DGRP lines, and estimated genetic (line) variances and covariances among these traits. To measure viral titre in the DGRP, we infected 5 vials of 10 flies for each line across 5 days, with a vial from each line being represented each day. After 8 DPI, living flies were killed and homogenised in Trizol for nucleic acid extraction and qPCR. To measure mortality following KV infection in the DGRP, we injected 3 vials of 10 flies of each sex and recorded mortality on alternate days until half the flies in the vial were dead (i.e. median survival time). Flies were transferred to fresh agar vials every 10 days. Mortality occurring in the first 3 DPI was assumed to be caused by the injection procedure and was removed from the analysis.

We fitted a multi-response linear mixed model in MCMCglmm to estimate heritability and genetic covariances among lines

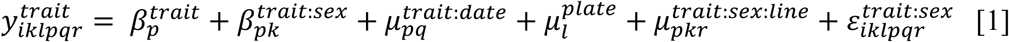

where 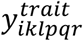 is the log-transformed relative viral titre or the duration until median mortality (LT50). We only estimated sex-specific fixed effects (*β^trait:sex^*) for LT50, because we did not measure titre in both sexes. The first part of the random effects model accounts for block effects due to date of injection (*μ*^*trait:date*^) and qPCR plate (*μ*^*plate*^). We assumed a 2×2 identity matrix as the covariance structure for *μ*^*trait:date*^, with effects associated with each trait from independent normal distributions. Effects for the *l*^*th*^ plate were assumed to be normally distributed. The second part of the random effects model (*μ*^*trait:sex:line*^) estimates the variance in each trait across lines, and wass allowed to vary by sex. We estimated all variance-covariance components of the 3×3 G matrix associated with *μ*^*trait:sex:line*^. Finally, we fitted separate error variances for each trait in each sex (*ε*^*trait:sex*^), where residuals were associated with independent normal distributions.

The diagonal elements of the *μ*^*trait:sex:line*^covariance matrix represent posterior distributions of genetic variances for viral titre in females, LT50 in females, and LT50 in males 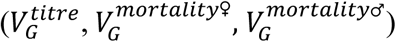. We calculated broad-sense heritability (i.e. line effects) for each trait as 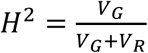, where *V_R_* is the residual variance associated with each trait, estimated in the model as *ε*^*trait:sex*^. However, heritabilities cannot readily be compared because of their dependence on the residual variance, which can be vastly different for different phenotypes (Houle, 1992). Therefore, we also calculated the coefficient of genetic variation (CV_G_) as 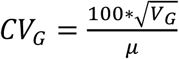, where V_G_ is standardised by the phenotypic mean (*μ*) and is more appropriate for comparisons across phenotypes. All confidence intervals reported are 95% highest posterior density intervals.

### Genome-wide association studies

We used measurements of viral titre and mortality following KV infection in the DGRP lines to perform a series of genome-wide association studies (GWAS). Although our power to detect small-effect genetic variants with only 125 lines is very low, past studies have demonstrated genetic variation in natural viral resistance in *Drosophila* is often dominated by few large effect variants (Contamine et al, 1989; Magwire et al, 2011; Magwire et al, 2012; Cogni et al, 2017; but see King and Long, 2017). We performed a GWAS on each phenotype separately by fitting an individual linear model for each variant in the genome using the full data. For the titre GWAS, we included focal SNP, qPCR plate, and date of injection as linear predictors. For the mortality GWAS, we included focal SNP, sex, and a sex-by-SNP interaction as linear predictors. Models were fitted using the base R linear model function ‘lm()’. We tested the significance of the SNP and SNP-by-sex predictors with a t-test, and we obtained significance thresholds for each GWAS by permuting genotypes across phenotypes 1000 times and recording the lowest *p*-value for each pseudo-dataset. Code for this model is made publicly available on figshare.

### Confirmation of GWAS hits

We chose 19 genes identified near significant GWAS hits to further test their involvement in KV infection. For each gene, we crossed a transgenic line containing a homologous foldback hairpin under the control of the Upstream Activating Sequence (UAS) to two GAL4 lines: *w*; P{UAS-3xFLAG.dCas9.VPR}attP40, P{tubP-GAL80^ts^}10; P{tubP-GAL4}LL7/TM6B, Tb^1^* (Bloomington line #67065; hereafter referred to as *tub-GAL4*) and *w*; P{GawB}Myo31DF^NP0001^/CyO; P{UAS-3xFLAG.dCas9.VPR}attP2, P{tubP-GAL80^ts^}2*

(Bloomington line #67067; hereafter referred to as *myo31DF-GAL4*). These lines drive GAL4 expression in the entire fly and in the gut, respectively, and contain a temperature-sensitive Gal80, which is able to inhibit GAL4 at the permissive temperature (18 degrees). RNAi lines included the following genes (BDSC numbers): *Pkcdelta* (28355), *btd* (29453), dos (31766), *tll* (34329), *Atg10* (40859), *Dgk* (41944), *Cip4* (53321), *hppy* (53884), *LpR2* (54461), *CG5002* (55359), *sev* (55866), *eya* (57314), *Gprk2* (57316), *Sox21b* (60120), *CG11570* (65014), *ATPCL* (65175), *Pdcd4* (66341), *CG7248* (67231), and *yin* (67334). As a control, we crossed the genetic background of the RNAi lines (Bloomington line #36304) to the two GAL4 lines. All crosses were made at 18 degrees. After eclosion, offspring were transferred to agar vials (10 flies per vial) at the non-permissive temperature (29 degrees) for two days to facilitate silencing of candidate genes, then injected with KV. We measured titre 5 DPI for 5 vials of each KV-infected genotype for each GAL4 driver. We used a linear mixed model to analyse log-transformed viral titre in each knockdown relative to the genetic background controls, with GAL4 driver as a fixed effect, gene knockdown as a random effect, and with separate error variances for each GAL4 driver. If the random effect associated with a candidate gene was significantly different from zero, we concluded this gene played a role in determining the outcome of infection by KV. The specification of gene as a ‘random effect’ allows comparison of each knockdown to all other knockdowns, accounting for any possible overall effect of overexpressing a dsRNA hairpin. As a proof of principle, we confirmed knock-down of the largest-effect gene (*Cip4*) using the DRSC FlyPrimerBank qPCR primers *Cip4_PP33370F* (ATTGCGGGAGTGACGCTTC) and *Cip4_PP33370R* (CTGTGTGGTGAGGTTCTGCTG). We did not assess knockdown efficiency for the other crosses, and any negative findings should be treated with caution.

### Sample preparation for RNA-sequencing

We next aimed to characterise the host expression response to KV infection, and whether this differed between males and females. We injected 6 vials of 10 flies for each sex with either the control gradient solution or with KV. After 3 DPI, we homogenised flies in Trizol, extracted total nucleic acid, and enriched the sample for mRNA through DNAse treatment and poly-A selection. We used the NEB Next Ultra Directional RNA Library Prep Kit to make strand-specific paired-end libraries for each sample, following manufacturer’s instructions. Libraries were pooled and sequenced by Edinburgh Genomics (Edinburgh, UK) using three lanes of an Illumina HiSeq 4000 platform with strand-specific 75 nucleotide paired end reads. We subsequently identified a low level of Drosophila A Virus contamination in both KV treated and untreated flies, reflecting the widespread occurrence of this virus in fly stocks and cell cultures. All reads have been submitted to the European Nucleotide Archive under project accession ERP023609.

### Differential expression analysis

We removed known sequence contaminants (primer and adapter sequences) from the paired end reads with cutadapt (V1.8.1; Martin, 2011) and mapped remaining reads to the *D. melanogaster* genome (FlyBase release r6.15) and to all known *Drosophila* virus genomes using STAR (V2.5.3a; Dobin et al, 2013), with a maximum intron size of 100 KB, but otherwise default settings. We counted the number of reads mapping to each gene using the featurecounts command in the subread package (V1.5.2; Liao et al, 2013) and used these raw count data as input to DESeq2 (V1.16.0; Love et al, 2014) for differential expression analysis. DESeq2 fits a generalised linear model for each gene, where read counts are modelled as a negative binomially distributed variable (Anders and Huber, 2010; Love et al, 2014) and includes a sample-specific size factor and a dispersion parameter that depends on the shared variance of read counts for genes expressed at similar levels (Anders and Huber, 2010; Love et al, 2014). Our design matrix included sex, KV infection status, and the interaction between the two, allowing us to test for expression changes following KV infection and how these changes differ between the sexes. To account for the unintended presence of DAV, and differences in the level of DAV within and between the treatments, we also include the relative titre of DAV as a continuous predictor. Using this model, we calculated log_2_ fold changes in DESeq2, and tested for significance using Wald tests. We used the ‘plotPCA’ function implemented in DESeq2 to perform principal component analysis of the rlog-transformed read count data (Love et al, 2014).

### GO term analysis

We performed five independent gene ontology (GO) term enrichment analysis, using: (1) genes with significant SNPs in the GWAS for titre; (2) genes with significant SNPs in the GWAS for mortality, (3) genes upregulated in either sex (p < 0.001); (4) genes downregulated in either sex (p < 0.001); and (5), genes significantly different between males and females (p < 0.05). For each of these gene lists, we tested for GO term enrichment using the ‘goseq’ R package (V1.26.0; Young et al, 2010), which accounts for the difference in power for detecting differential expression caused by gene length, and tests for significant over-representation of genes in a GO term.

## Results and Discussion

### Isolation of Kallithea Virus

We isolated Kallithea Virus (KV) by gradient centrifugation following 4 rounds of serial passage in flies. Many laboratory fly stocks and cell culture lines are persistently infected with RNA viruses (Brun and Plus, 1980; Webster et al, 2015), and following serial passage we identified co-infections of Drosophila A Virus (DAV), Nora Virus, and Drosophila C Virus (DCV) by PCR. The high prevalence of these viruses in laboratory stocks presents a substantial hurdle in the isolation of new *Drosophila* viruses, requiring the separation of the new viruses of interest. Although this can be relatively simple (e.g. separating enveloped from non-enveloped viruses), most of the recently identified *Drosophila* viruses (Webster et al, 2015; Webster et al, 2016; Medd et al, 2017) are from ssRNA virus families with buoyant densities similar to common laboratory infections. To exclude these from our isolate, we concentrated KV using a 1.18 g/mL cushion, retaining KV at the interphase, but excluding most of the contaminating RNA viruses. Subsequent equilibrium density gradient centrifugation produced a KV band at 1.17 g/mL, and with some DAV contamination at approximately 1.20 g/mL (Figure 1A). Although nudiviruses have not previously been prepared using an iodixanol gradient, the equilibrium buoyant density was consistent with the lower buoyant densities of enveloped particles (Feng et al, 2013) and similar to other enveloped dsDNA viruses (e.g. Herpesviruses: 1.15 g/mL). KV was estimated to be an approximately 650-fold higher concentration than DAV at 1.17 g/mL, and we were unable to identify intact DAV particles by electron microscopy (KV shown in Figure 1B). KV is morphologically similar to Oryctes rhinoceros nudivirus (Huger, 2005), with an enveloped rod-shaped virion approximately 200 nm long and 50 nm wide.

**Figure 1:**
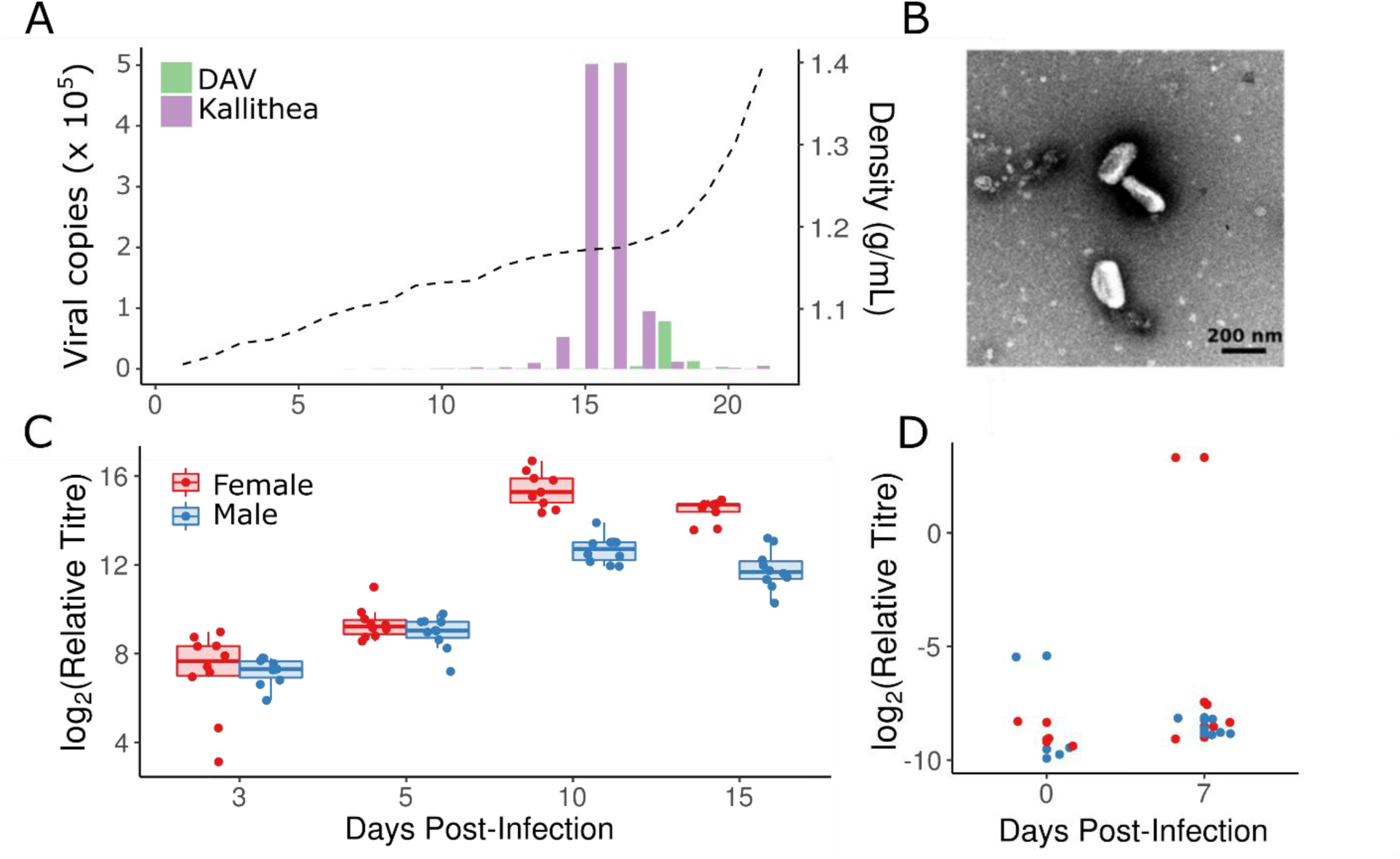
Isolation of KV and growth in flies. (A) Density gradient and virus titre: Kallithea virus (purple) was effectively separated from DAV (green) at 1.18 g/mL (dotted line) in fractions 15 and 16 of an iodixanol gradient. (B) Transmission electron micrograph of KV-positive fractions showed KV to be a rod-shaped enveloped particle, as has been described previously for other nudiviruses (Huger, 2005). We did not observe unenveloped KV particles, bacteria, or RNA viruses in the isolate. (C) Relative viral titres normalised by the number of fly genomic copies and virus levels at time zero in each sex. Each point represents a vial of 10 flies. Viral titres peaked at 10 days post-infection, and were generally higher in females (red) than males (blue) late in infection. (D) We were able to infect adult flies orally by applying the viral isolate to *Drosophila* medium, although relative copy number of the virus was very low and infection was inefficient, with only 2 of 16 vials (each of 10 flies) having increased titre after one week, indicating an infectious rate lower bound of ∼1% at 5×10^3^ ID50.

### Kallithea virus growth in flies

We injected the KV isolate into *Drosophila Oregon R* males and females, with and without *Wolbachia*, and measured viral titre at four time-points by qPCR. In females, KV increased approximately 45,000-fold by day 10, and then began to decrease (Figure 1C). In males, the KV growth pattern was altered, growing more slowly (or possibly peaking at an earlier un-sampled time point), resulting in a 7-fold lower titre than in females after 10-15 days, (nominal MCMC p-value derived from posterior samples, MCMCp= 0.002). *Wolbachia* did not affect virus growth rate in either sex (MCMCp = 0.552, Figure S1), reaffirming previous findings that *Wolbachia* do not offer the same protection against DNA viruses in *Drosophila* as they do against RNA viruses (Teixeira et al, 2008).

Nudiviruses have previously been reported to spread through sexual and faecal-oral transmission routes. The *Drosophila innubila* Nudivirus (DiNV), a close relative of KV, is thought to spread faecal-orally, so we tested whether KV can spread through infected food. We found that although oral transmission occurred, it was relatively inefficient (Figure 1D). However, the concentration of DiNV found in *D. innubila* faeces is broadly similar to our KV isolate after gradient centrifugation (Unckless 2011; Figure 1D), but the administered suspension had been diluted 50-fold and may consequently provide a lower dose than flies encounter naturally. To explore the potential for transovarial vertical transmission or gonad-specific infections following sexual transmission (as reported for Helicoverpa nudivirus 2; Burand et al, 2012), we also performed qPCR on dissected ovaries and the remaining carcasses at 3 DPI (Figure S2). We found that KV was highly enriched in the carcass relative to the ovaries. Although intra-abdominal injection could influence KV tissue-specificity, there were still substantial levels of KV in the ovaries, indicating there is not a complete barrier to infection. These results imply that KV is likely transmitted faecal-orally, as are closely related nudiviruses, but explicit tests for transovarial or sexual transmission are required.

*Sex-specific mortality, lethargy, and altered fecundity patterns following KV infection Drosophila innubila* infected with DiNV suffer fitness costs including increased mortality and decreased fecundity (Unckless 2011). We investigated KV-induced mortality in *D. melanogaster* by injecting males and females, with and without *Wolbachia*, with either control gradient solution or KV. We found that KV caused slightly, but significantly, increased mortality in females compared with controls (21% dead by day 20, vs. 11% in controls, MCMCp = 0.001), but caused a dramatically increased mortality in males compared to females (63% dead by day 20, vs. 14% in controls, sex:virus interaction MCMCp < 0.0001; Figure 2A). Therefore, males appear less tolerant of infection by KV, displaying increased mortality and a lower titre than females. We confirmed the KV-induced male death was not caused by DAV or other unknown small unenveloped RNA viruses present in our initial isolate, as chloroform treatment of the KV isolate eliminated treatment associated mortality (Figure S3). Male-specific costs of infection are widespread across animal hosts and their pathogens (e.g. Zuk, 2009), and reduced male tolerance has been found in flies infected with Drosophila C Virus (DCV) (Gupta and Vale, 2017). We found that *Wolbachia* infection had no detectable effect on KV-induced mortality in males or females, and thus does not affect tolerance (MCMCp = 0.20; Figure S1). This is consistent with previous studies showing that *Wolbachia* infection affects resistance and tolerance to RNA viruses but not a DNA virus (Teixeira et al, 2008).

**Figure 2:**
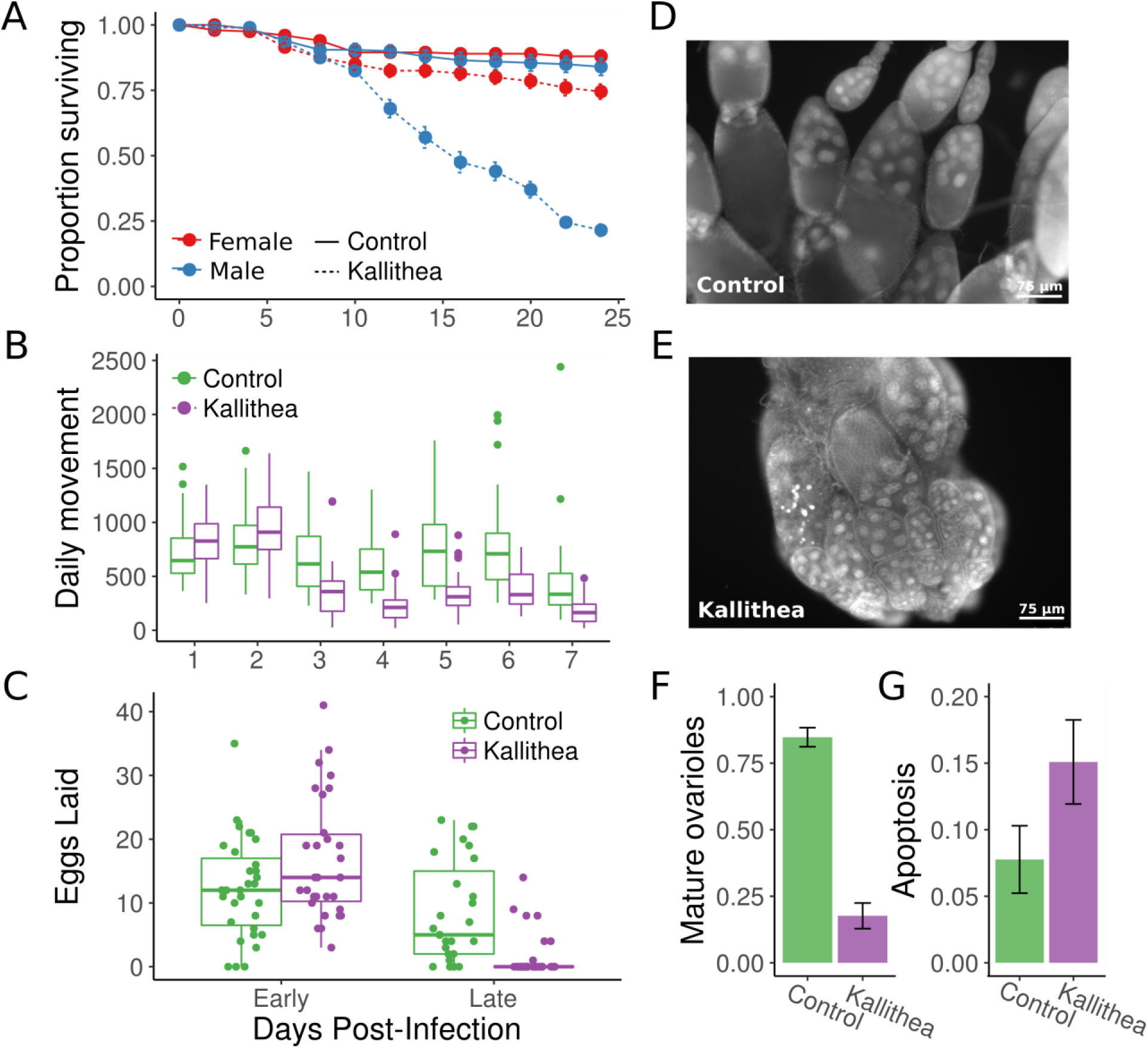
KV causes male-specific mortality, increased lethargy, and decreased fecundity. (A) Injection of KV virus into *OreR* flies led to sex-specific mortality. Infected females (red dotted line) experienced a small but significant increase in mortality, but males (blue dotted line) experienced a significantly larger rate of mortality after day 10. Flies injected with control gradient solution were unaffected (solid lines). Each point is the mean and standard error for the proportion of flies alive in each vial (10 vials of 10 flies). (B) Although females remained alive, they were more lethargic. We assessed daily movement of flies injected with either chloroform-inactivated KV (green) or active KV (purple). KV-infected flies moved less from days 3-7 post-infection. (C) Females also displayed altered egg laying behaviour. Thirty pairs of flies were injected with inactive chloroform treated KV (green) or active KV (purple). KV-infected flies laid a slightly higher number of eggs during early infection (1 and 2 DPI) but laid significantly fewer eggs in late infection (7 and 8 DPI). This reduction in egg laying is due to a shutdown of oogenesis before vitellogenesis (D, E), and ovaries from KV-infected flies house a lower proportion of ovarioles that include late-stage and mature egg chambers (F) and a higher proportion which contain apoptotic nurse cells (G). Ovaries were analysed 10 DPI.

We next tested whether female flies suffer sub-lethal fitness costs, by monitoring fly movement for a week following infection. KV-infected female flies showed similar movement patterns to chloroform-treated KV-injected flies for two days post-infection, but from three days post-infection moved significantly less (∼70% reduction relative to controls; MCMCp < 0.001; Figure 2B). We conclude that females suffer from increased lethargy resulting from KV infection. In a natural setting, this could translate into fitness costs associated with increased predation, and reduced egg dispersal, mating, and foraging.

Finally, we tested whether KV infection resulted in decreased fecundity by monitoring the number of eggs laid for 12 days post-infection. We found that infected females exhibited markedly different egg laying patterns (MCMCp < 0.001; Figure 2), with KV-infected flies consistently laying fewer eggs at 7 and 9 days post-inoculation. This reduction in egg-laying during late infection could be due to a behavioural response or a cessation of oogenesis. To differentiate between these possibilities, we dissected ovaries, and determined the proportion of ovarioles that contained mature egg chambers. We found that ovaries from KV-infected flies halt oogenesis around stage 8 (MCMCp < 0.001), before vitellogenesis, and house an increased number of apoptotic egg chambers (MCMCp < 0.001) (Figure 2). This phenotype is similar to that seen upon starvation (Jouandin et al, 2014), and could be the manifestation of a trade-off to reroute resources to fighting infection, or of sickness-induced anorexia (e.g. Ayres and Schneider, 2009). Alternatively, this could be a direct consequence of viral infection, consistent with the gonadal atrophy reported for HzNV-2 (Burand et al, 2012). Future studies should address whether this phenotype is a direct or indirect consequence of infection, and if the latter, whether it is orchestrated by the host or the virus.

### Variation in titre and mortality following KV infection

The DGRP (Mackay et al, 2012) have previously been used to dissect genetic variation underlying resistance and tolerance to bacterial, fungal, and viral pathogens (Magwire et al, 2012; Bou Sleiman et al, 2015; Wang et al, 2017; Howick and Lazzaro, 2017). We infected 125 DGRP lines with KV and estimated broad-sense heritabilities (H^2^: the proportion of phenotypic variance attributable to genetic line) and coefficients of genetic variation (CV_G_: a mean-standardised measure of genetic variation) in viral titre and LT50 values in females, and LT50 values in males (Table 1). Our estimates of H^2^ and CV_G_ fall within the range found for resistance to other pathogens in the DGRP, although direct comparison is difficult as studies are inconsistent in the statistics used to report genetic variation. H^2^ in survival following infection with an opportunistic bacterium or fungus was similar to our estimate for survival following KV infection (*Pseudomonas aeruginosa*: H^2^ in males = 0.47, H^2^ in females = 0.38; *Metarhizium anisopliae*: H^2^ in males = 0.23, H^2^ in females = 0.27; Wang et al, 2017), although comparing heritability can be easily confounded by differences in environmental (residual) variance (Houle, 1992). Genetic variation in resistance has also been measured in response to two non-native *D. melanogaster* viruses (Flock House Virus and *Drosophila affinis* Sigma Virus) and two native viruses (DCV and DmelSV) in females of the DGRP. Of these, the lowest heritabilities are those associated with resistance to non-native fly viruses (FHV: h^2^ = 0.07, CV_G_ = 7; *D. affinis* sigma virus: h^2^ = 0.13), and the highest are associated with native fly viruses (DCV: h^2^ = 0.34, CV_G_ = 20; DmelSV: h^2^ = 0.29). Although Magwire et al (2012) inferred narrow sense heritability (h^2^) as half V_G_ and accounted for the homozygosity of inbred lines when inferring CV_G_, it is clear that V_G_ for resistance to KV is closer to the V_G_ for resistance to other native fly viruses than to non-native ones, at least for survival. It is also notable that CV_G_ estimates for survival are higher than estimates for titre, consistent with the observation that traits more closely related to fitness are expected to have higher CV_G_ values (Houle, 1992).

**Table 1:** Trait means, genetic variance (V_G_), total phenotypic variance (V_P_), heritability (H^2^), and coefficient of genetic variation (CV_G_) in titre and mortality following KV.

| **Trait** | **Mean** | **V<sub>G</sub>** | **V<sub>P</sub>** | **H<sup>2</sup>** | **CV<sub>G</sub>** |
| --- | --- | --- | --- | --- | --- |
| *Titre* | 1.96 [1.31 - 2.63] | 0.05 [0.03 - 0.09] | 0.28 [0.24 - 0.32] | 0.19 [0.1 - 0.29] | 11.8 [8.4 - 15.2] |
| *LT50* ♀ | 25.8 [22.6 - 29.0] | 25.2 [17.4 - 33.8] | 44 [35.7 - 52.6] | 0.57 [0.47 - 0.67] | 19.5 [16.2 - 22.6] |
| *LT50* ♂ | 19.6 [16.4 - 22.7] | 10.9 [5.9 - 16.5] | 35.7 [30.1 - 41.7] | 0.3 [0.18 - 0.42] | 16.7 [12.7 - 21] |

We calculated genetic correlations between male and female mortality, and between viral titre and mortality in females (Figure 3). Note that we found no correlation between survival time following KV infection and published estimates of longevity in the absence of infection, nor to resistance to any other RNA viruses (Ivanov et al, 2015; Magwire et al, 2012). We found a strong positive correlation between males and females in median survival time following KV infection (0.57 [0.34-0.78]; MCMCp <0.001), such that lines in which infected males die quickly are also lines in which infected females die quickly, suggesting a shared genetic basis for early lethality following infection. We also surprisingly find a positive genetic correlation between viral titre and LT50 values (r = 0.32 [0.05-0.59], MCMCp = 0.017), such that fly lines that achieved higher titres on day 8 tended to live slightly longer. However, the effect size is small (a doubling of viral titre led to a half-day increase in median survival time) and the result is only marginally significant. The absence of a negative correlation is counter-intuitive, and contrasts with infection of the DGRP with *Providencia rettgeri* and *Metarhizium anisopliae*, and infection across *Drosophila* species with DCV, where fly lines or species with higher parasite loads suffer increased mortality (Longdon et al, 2015; Wang et al, 2017; Howick and Lazzaro, 2017). This apparent decoupling of titre and mortality could result from inherent costs associated with the induction of an immune response, whereby flies that raise a more potent immune response keep KV at lower titres but induce greater tissue damage.

**Figure 3:**
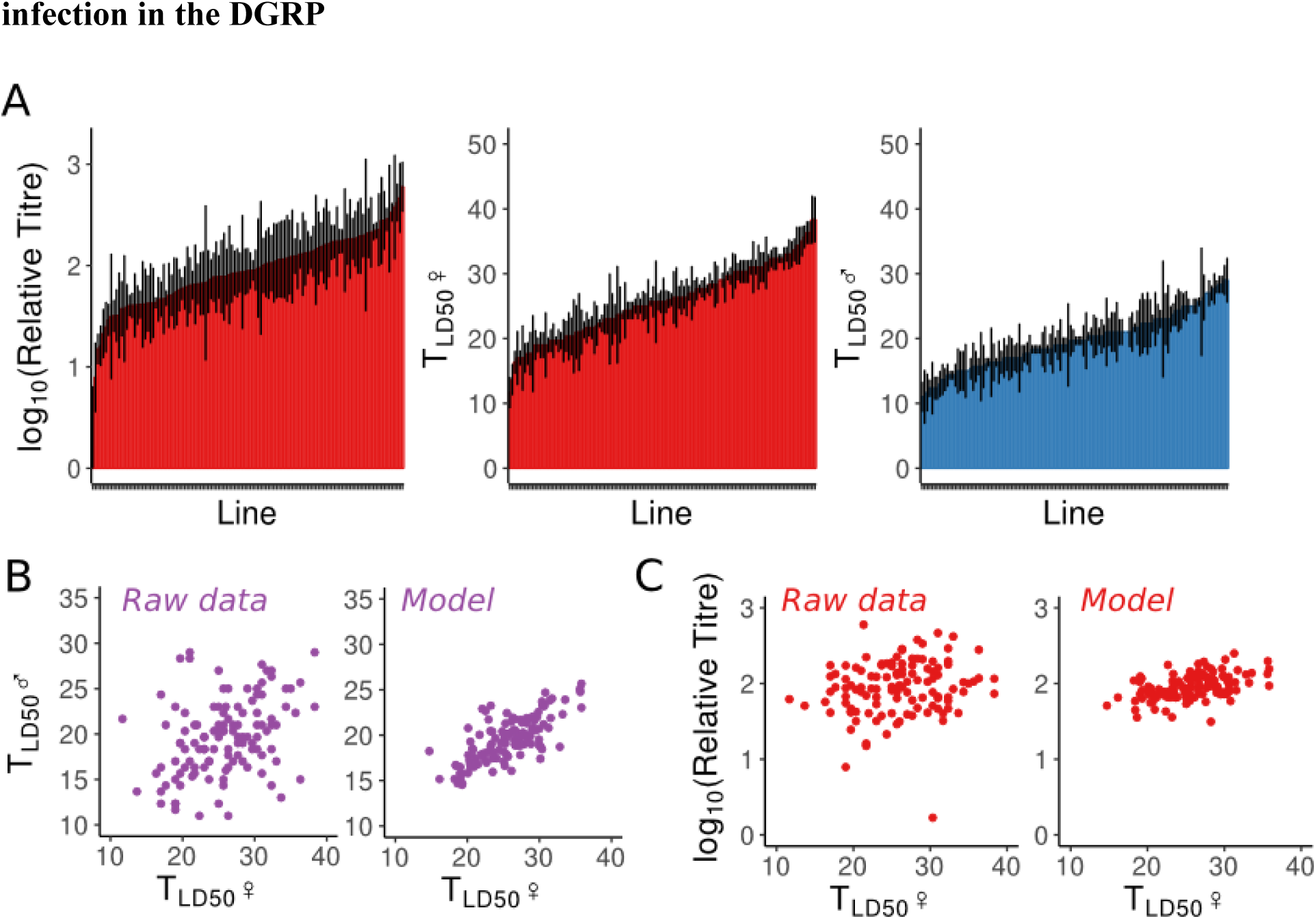
Genetic variation in resistance to KV. (A) We measured LT50 in both sexes, and titre in females, following KV injection in the DGRP. For titre, each bar represents the mean (and standard error) titre relative to fly genome copy-number, as assessed by qPCR for 5 vials of 10 flies for each of 125 DGRP lines. For LT50, each bar represents the mean time until half the flies (in a vial of 10) were dead, for three vials per line, per sex. (B, C) We used a multi-response linear mixed model to calculate genetic correlation between the traits. Shown are the raw data (left), and the estimated line effects (right) after accounting for any injection date and qPCR plate effects, and for estimated variance among lines. Each point is a DGRP line measured for both phenotypes. We find a strong positive correlation between male and female LT50 values (B). We also observe a weak positive correlation between titre and LT50 (C).

### Identification of candidate genes underlying host variation in KV titre

Using the phenotypes in the DGRP lines measured above, we performed a genome-wide association study to identify candidate genes underlying variation in titre, LT50, and differences in LT50 between the sexes. We found 10 SNPs (9 near genes) that were significantly associated with viral titre (p_ran_ < 0.05, based on 1000 permutations of phenotypes across lines; Figure 4). The SNP with the smallest p-value appeared in *Lipophorin receptor 2* (*LpR2*), which encodes a low-density lipoprotein receptor, previously found to be broadly required for flavivirus and rhabdovirus cell entry (e.g. Agnello et al, 1999; Albecka et al, 2012; Finkelshtein et al, 2013).

**Figure 4:**
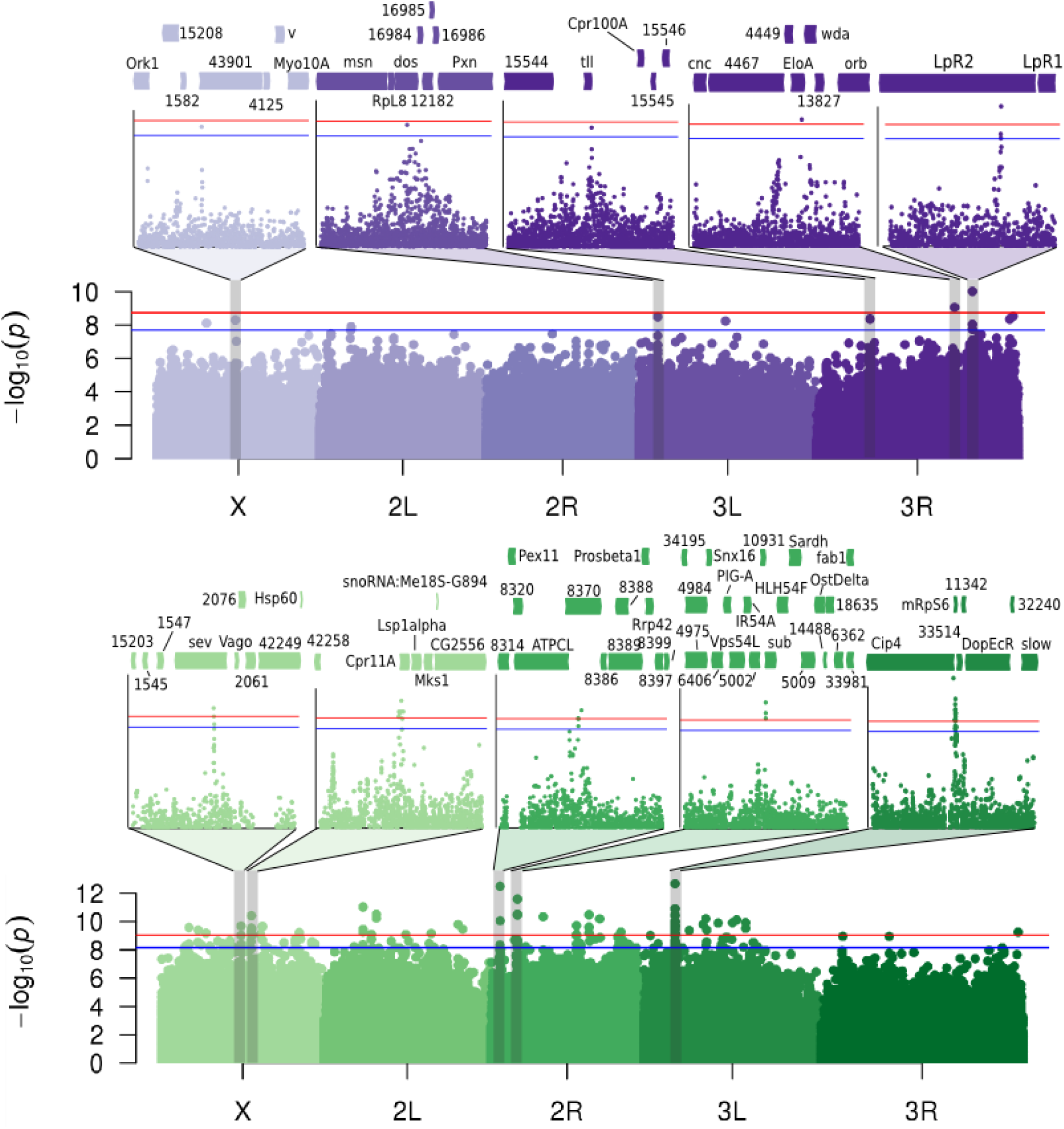
Genome-wide association of polymorphism in the DGRP with KV-induced titre and mortality. Manhattan plots showing the p-value for the effect of each polymorphism on viral titre (purple) and mortality (green). The top SNPs for each phenotype are shown in expanded inset panels, including surrounding genes. For clarity “CG” is omitted from gene identifiers. Horizontal lines show significance thresholds obtained through randomisation (p_ran_ = 0.05 in blue; p_ran_ = 0.01 in red).

We tested whether these candidate polymorphisms were enriched in any molecular, biological, or cellular processes using a GO enrichment analysis, and found the top hit to be the torso signalling pathway with 2 genes of 34 in the category (p = 0.0004), *tailless* and *daughter of sevenless* (*dos*). Torso signalling is upstream of extracellular-signal-regulated kinase (ERK) pathway activation in some tissues, and human orthologues of *dos* (GAB1/GAB2/GAB4) are cleaved by an enterovirus-encoded protease, thereby activating ERK signalling and promoting viral replication (Deng et al 2015; Deng et al, 2017). ERK signalling is also an important regulator of virus replication in the fly midgut, where it couples nutrient availability with antiviral activity (Xu et al, 2013; Liu et al, 2017). See Table S1 for a list of all nominally significant SNPs with associated locations, mutation types (e.g. intronic, synonymous coding, etc), nearby genes, p-values, effect sizes, and GO terms.

### Identification of candidate genes underlying host variation in KV-induced mortality

We found 86 SNPs (65 near genes) that were significantly associated with LT50 following KV infection in the DGRP (p_ran_ < 0.05; Figure 4, Table S1), none of which were identified in the GWAS for viral titre. We performed a GO enrichment analysis, and found genes associated with these SNPs were enriched for hydrolase activity (top molecular function GO term, p = 0.0004), stem cell fate determination (top biological process GO term, p = 0.002), and in the plasma membrane (top cell component GO term, p = 0.004), among others (Table S2). Of these 86 SNPs, we found 34 (26 near genes) that were highly significant, and selected these for further analysis and confirmation (p_ran_ < 0.01; see Table S1 for all significant SNPs in). The polymorphism with the most confident association was located in *Cdc42-interacting protein 4* (*Cip4*), a gene involved in membrane remodelling and endocytosis (Leibfried et al, 2008; Fricke et al, 2009). This SNP is intronic in the majority of *Cip4* transcripts, but represents a nonsynonymous polymorphism segregating leucine and proline in the first exon of *Cip4-PB* and *Cip4-PD* isoforms, perhaps indicating spliceform-specific effects on KV-induced mortality. Of particular interest from the remaining 33 highly significant SNPs was a synonymous SNP in the receptor tyrosine kinase, *sevenless*, known to interact with *dos* (above), and seven genes (*Dgk*, *Atg10, ATPCL, Hppy, Pkcdelta, Gprk2, Pdcd4*) previously implicated in viral pathogenesis or general immune processes. Of these, three (*Gprk4, hppy, Pkcdelta*) are involved in NFκB signalling (Valanne et al, 2011, Chuang et al, 2011; Loegering and Lennartz, 2011). *ATPCL* was identified in an RNAi screen for factors regulating Chikungunya virus replication in humans (Karlas et al, 2016) and is involved in the late replication complexes of Semliki Forest Virus (Varjak et al, 2013). Finally, *Atg10* and *Pdcd4* are involved in autophagy and apoptosis, respectively, both broadly antiviral cellular functions known to have a role in antiviral immunity in *Drosophila* (Shelly et al, 2009; Lamiable and Imler, 2014). We found no SNPs significantly associated with sex-specific KV-induced mortality (Figure S4).

### Confirmation of GWAS hits

We chose 18 GWAS-candidate genes with available UAS-driven RNAi constructs to verify their involvement in KV infection. We found that knockdown of *Cip4* and *CG12821* caused significantly increased viral titre, and knockdown of *sev* and *dos* resulted in significantly decreased viral titre, relative to other knockdown lines (Figure 5; Figure S5). We confirmed *tub-GAL4>Cip4^IR^* flies had reduced (26% of wild-type) *Cip4* RNA levels and a concomitant increase in viral titre relative to the genetic background control (3.4-fold increase, 95% C.I. 1.3 – 9.6 fold) (Figure 5). This confirms that *Cip4* is a KV restriction factor that likely segregates functional polymorphism affecting survival following KV infection (Figure 5). It is known that baculovirus budded virions enter cells through clathrin-mediated endocytosis or micropinocytosis (Long et al, 2006; Kataoka et al, 2012), and gain their envelope at the cell membrane upon exit (Blissard and Rohrmann, 1990). Cip4 could therefore plausibly enact an antiviral effect by limiting KV cell entry or spread, perhaps through its known function in cell membrane remodelling and trafficking.

**Figure 5:**
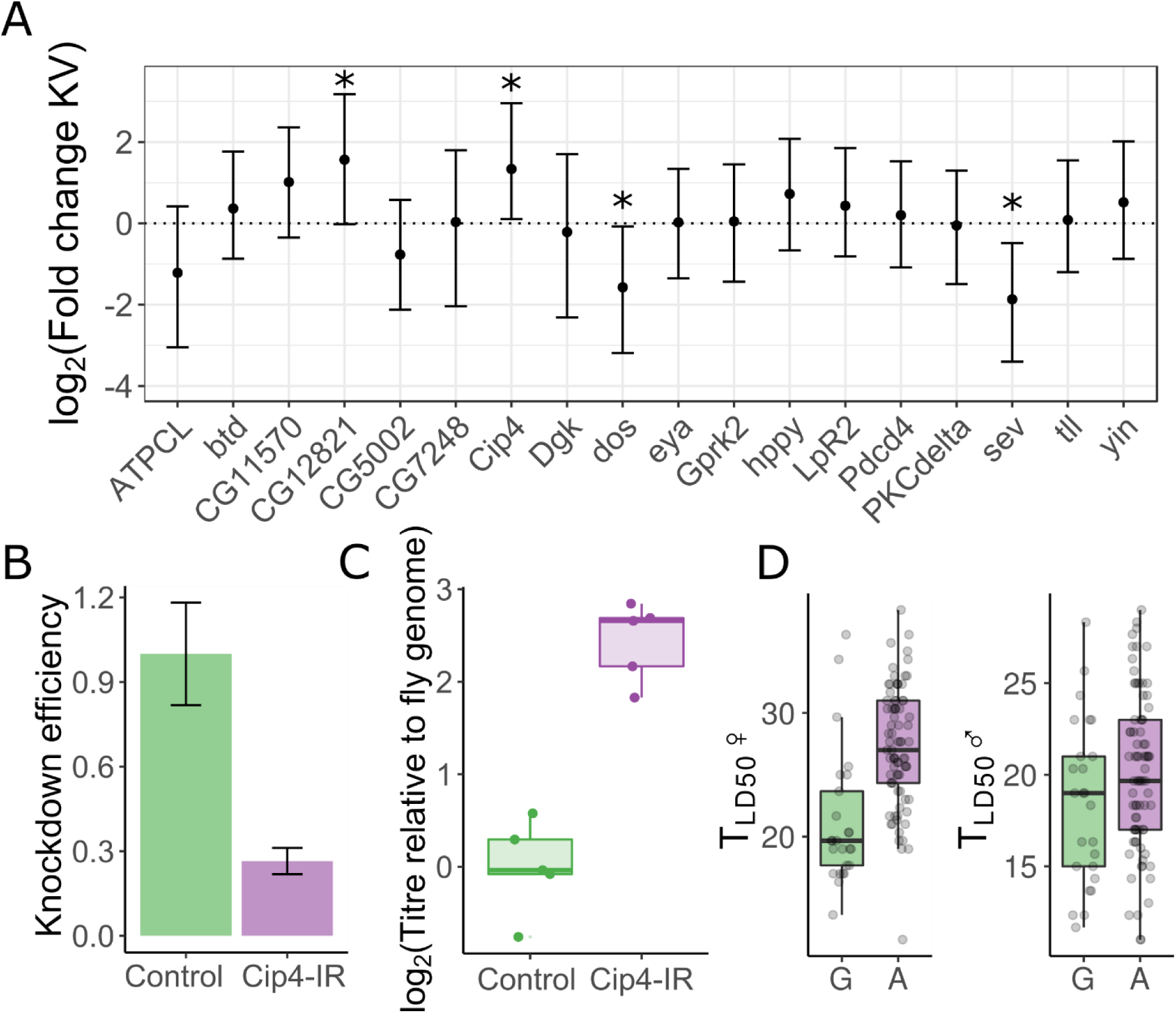
Confirmation of antiviral genes identified in GWAS. KV titre was measured in flies expressing a foldback hairpin targeting 18 genes identified in the GWAS, using GAL4 lines that knock each down in either the whole fly or specifically in the gut. (A) The data were used to estimate random effects associated with each gene knock down, plotted with 95% highest posterior density intervals. (B) Knock-down of the most confident association in the GWAS, *Cip4*, caused reduced *Cip4* RNA levels and (C) increased viral titre. (D) The associated variant (3L_4363810_SNP), was polymorphic (G/A), representing a nonsynonymous polymorphism in some splice variants, and survival following KV infection was significantly increased in fly lines with the “A” genotype, especially in females. Each point in comparison of survival in the two genotypes is a line mean. (*MCMCp < 0.05)

### Differential expression following KV infection

Previous transcriptional profiling in response to RNA virus infection has shown upregulation of heatshock proteins, JAK-STAT, JNK, and Imd pathways (Dostert et al, 2005; Zhu et al, 2013; Kemp et al, 2013; Merkling et al, 2015). However, the *D. melanogaster* expression response to a DNA virus has not previously been investigated. We separately injected male and female flies with control gradient solution or KV and extracted mRNA for sequencing 3 days post-infection. KV gene expression increased dramatically 3 days-post inoculation, consistent with our qPCR analysis of genome copy-number (Figure 1, Figure S9). Although not previously detectable by PCR, RNAseq read mapping identified a low level of DAV in both control and KV-infected flies, with an overall higher level in KV-infected flies. To account for this potentially confounding contaminant, we fitted the number of DAV-mapped reads as a covariate in the differential expression analysis, and used a stringent Benjamini–Hochberg adjusted significance threshold of *p* < 0.001 to infer nominal significance. We found 54 genes upregulated and 79 genes downregulated in response to KV in either males or females (Figure 6; Table S3). There was no enrichment for GWAS hits among the KV-responsive genes (Figure S6). Principal components analysis on depth-normalised read counts separated males and females along PC1 and partially separated KV-infected and control-injected libraries along PC3 (Figure S7). GO term analysis identified ‘defense response to virus’ (p = 3.1×10^−4^), ‘serine peptidase activity’ (p = 1.2×10^−7^, identified in part due to downregulation of Jonah family serine proteases), and ‘chorion’ (p < 1×10^−8^) as the most highly enriched biological process, molecular function, and cellular component, respectively (Figure 6; Table S4).

**Figure 6:**
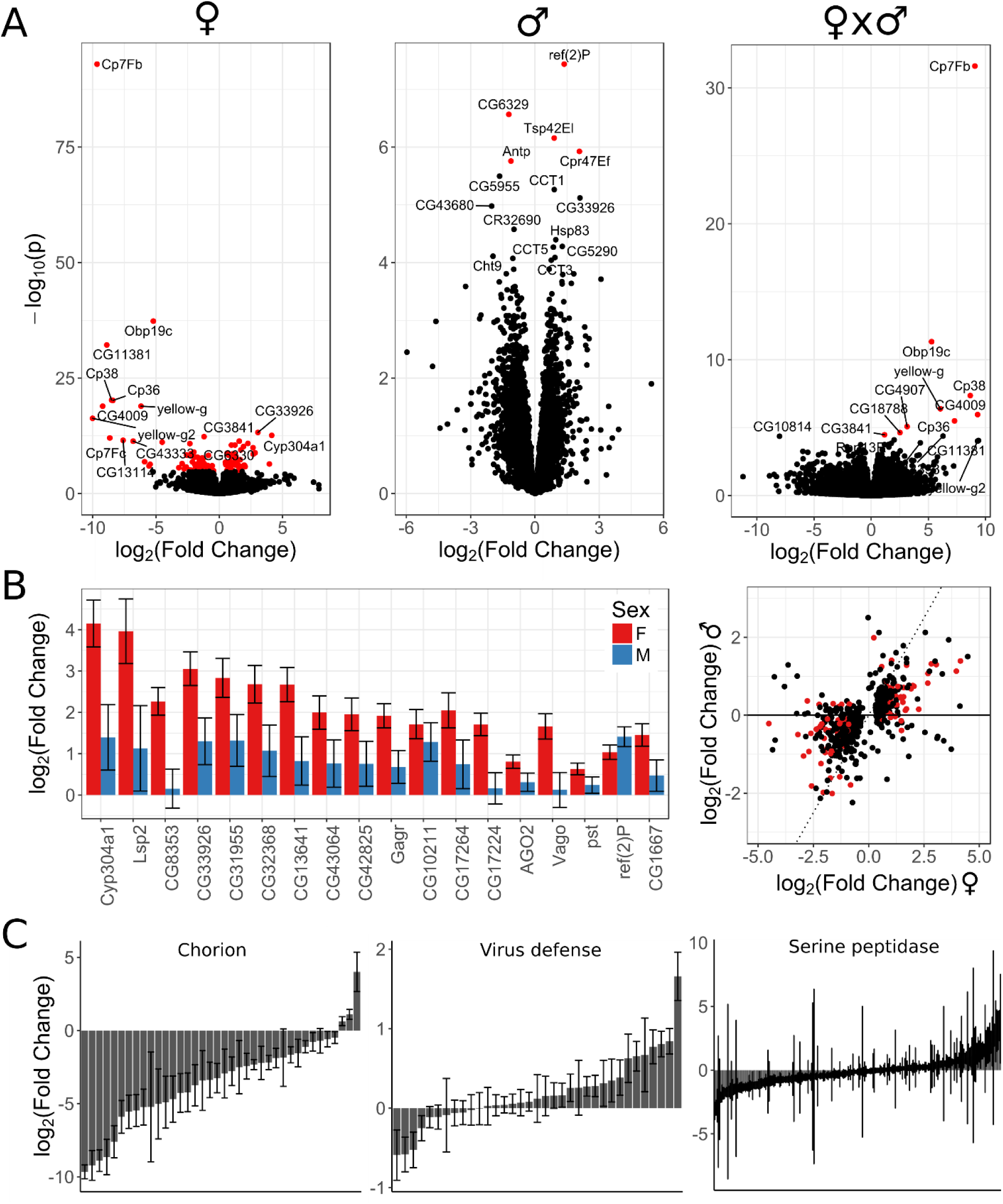
KV induces differential regulation of chorion, virus defense, and serine endopeptidase genes. (A) Volcano plots showing fold changes and p-values from Wald tests for differential expression of *D. melanogaster* genes following KV infection for females (left), males (center), those different between the sexes (right), with DAV read count fit as a covariate and nominal significance threshold of p < 0.001. In each panel, the genes with the smallest p-values are labelled. All of the genes significantly differentially regulated between the sexes are those highly significant in females. (B) Highly induced genes are mostly functionally unannotated, but include some with known roles in viral pathogenesis. (C) The male response is correlated to females, but muted, with few genes identified as significantly differentially expressed in males. Genes with weak evidence of differential expression in either sex (p < 0.05) are plotted, where the dotted line represents a perfect correlation, and red points are genes identified as significantly differentially expressed. (D) The top GO enrichment terms for each GO class (Molecular Function, Biological Process, Cellular Component) were genes involved the chorion, virus defense, and serine peptidase activity. For each plot, estimated fold changes and their associated standard errors are plotted for every gene in a GO term, regardless of the significance of the Wald test. Generally, chorion genes were downregulated, virus defense genes were upregulated, and serine peptidases were downregulated.

There are few described induced antiviral immune effectors in *Drosophila* (e.g. Lamiable and Imler, 2014). In line with this we observe 57% of differentially expressed genes have not yet been named (i.e. “CG” genes), significantly greater than the genome-wide rate of 41% (p = 3×10^−4^), and the most highly induced genes have not been implicated in viral pathogenesis. The cytochrome P450 family gene *Cyp304a1* was most highly upregulated, concomitant with the upregulation of four other genes in this family (*Cyp309a1, Cyp309a2, Cyp4p3*, and *Cyp6a20*). The next most highly induced genes include the hemocyanin *Larval serum protein 2*, the cytidine deaminase *CG8353*, four genes without functional annotation or recognisable domains (*CG33926*, *CG31955*, *CG32368*, *CG13641*), and an additional six genes without functional annotation (*CG43064, CG42825, Gagr, CG10211, CG17264*, and *CG17224* –the last two of which are adjacent on chromosome arm 2L). We also note the striking but variable upregulation of 11 of the 24 Tweedle genes (Figure S8) in some (but not all) of the infected samples. These are secreted, insect-specific cuticle proteins that regulate body shape (Guan et al, 2006), and are also upregulated in response to Sindbis virus infection in cell culture (Graveley et al, 2011, Pers Comm to Flybase), perhaps suggesting a general role in viral pathogenesis.

Genes with known involvement in viral pathogenesis were also found to be induced following KV infection. The RNAi effector *AGO2* was upregulated, consistent with the previous results that DNA viruses are a target of the RNAi pathway (Kemp et al, 2013; Bronkhorst et al, 2012; Bronkhorst et al, 2014). *Vago*, an antiviral factor downstream of Dicer-2 (Deddouche et al, 2008), was upregulated and was also adjacent to a SNP found in the mortality GWAS (*dos*; Figure 4, Figure 7), as were *pastrel* and *ref(2)P* induced, identified in previous genome wide association analyses for resistance to DCV and DMelSV, respectively. Finally, we found that KV induced expression of *CG1667*, the *Drosophila* homologue of STING. The vertebrate cGAS-STING pathway is involved in cytosolic DNA sensing and activation of immune factors in response to DNA virus infection (Chen et al, 2016), and the upregulation of *CG1667* may suggest that this is another pathway conserved between *Drosophila* and vertebrates.

As we had observed male and female differences in KV-induced mortality and titre (Figure 1, Figure 2), we tested for sex-specific transcriptional regulation in response to KV infection. We found that females and males had similar patterns of differential expression following KV infection (spearman rank correlation coefficient, ρ = 0.57, p = 2.2×10^−16^), although the male response was often less potent (Figure 6). Nine genes were significantly differentially expressed (p < 0.05) between the sexes specifically in response to KV (Table S3), and these were all downregulated in females and highly enriched for genes associated with the chorion (Figure 6, Table S4). Strikingly, all but three genes classified with the GO term ‘chorion’ were downregulated in females (Figure 6), consistent with the observed reduction in mature ovarioles and eggs during late infection, and implying a substantial reorganization of oogenesis (Figure 2). We did not identify any previously described immune genes with significant sex-specific regulation during KV infection, although *Vago* upregulation appeared superficially higher in females (Figure 6; Table S3).

## Conclusions

We have isolated Kallithea Virus, a dsDNA nudivirus that naturally infects *D. melanogaster*, and find it to be experimentally tractable. KV infection leads to reduced fertility and movement in females, highlighting the importance of measuring fitness associated traits besides longevity. Although males suffered greater mortality than females, they achieved lower titres, consistent with increased resistance and/or reduced tolerance in males. Similar to RNA viruses, we identified moderate host genetic variation in resistance to KV infection, however, we found that the underlying genetic architecture of this variation is unlike previously studied RNA viruses of *D. melanogaster*, in which a high proportion of genetic variation was apparently determined by a small number of loci. This could reflect a difference in the co-evolutionary dynamics between *D. melanogaster* and KV, versus other RNA viruses such as DCV and DMelSV. The *D. melanogaster* transcriptional response to KV included genes with known involvement in viral pathogenesis, but also genes that could represent infection responses distinctive to DNA viruses or KV, including downregulation of chorion genes. Upregulation of widely conserved immune factors, such as *STING*, represent promising candidates involved in fly antiviral immunity, and demonstrate the continued utility of the *Drosophila* system for understanding host-virus interactions.

## Acknowledgements

We thank John Pool for the original fly collection that contained the KV isolate; Megan Wallace, Steve Mitchell, and Naomi Letley for help with experimental work; Pedro Vale, Katy Monteith, and Fergal Waldron for creation, upkeep, and sharing of the outbred DGRP population; Craig Walling and Jarrod Hadfield for statistical advice, and members of the Obbard and Vale labs for discussion.

---

**Figure S1:**
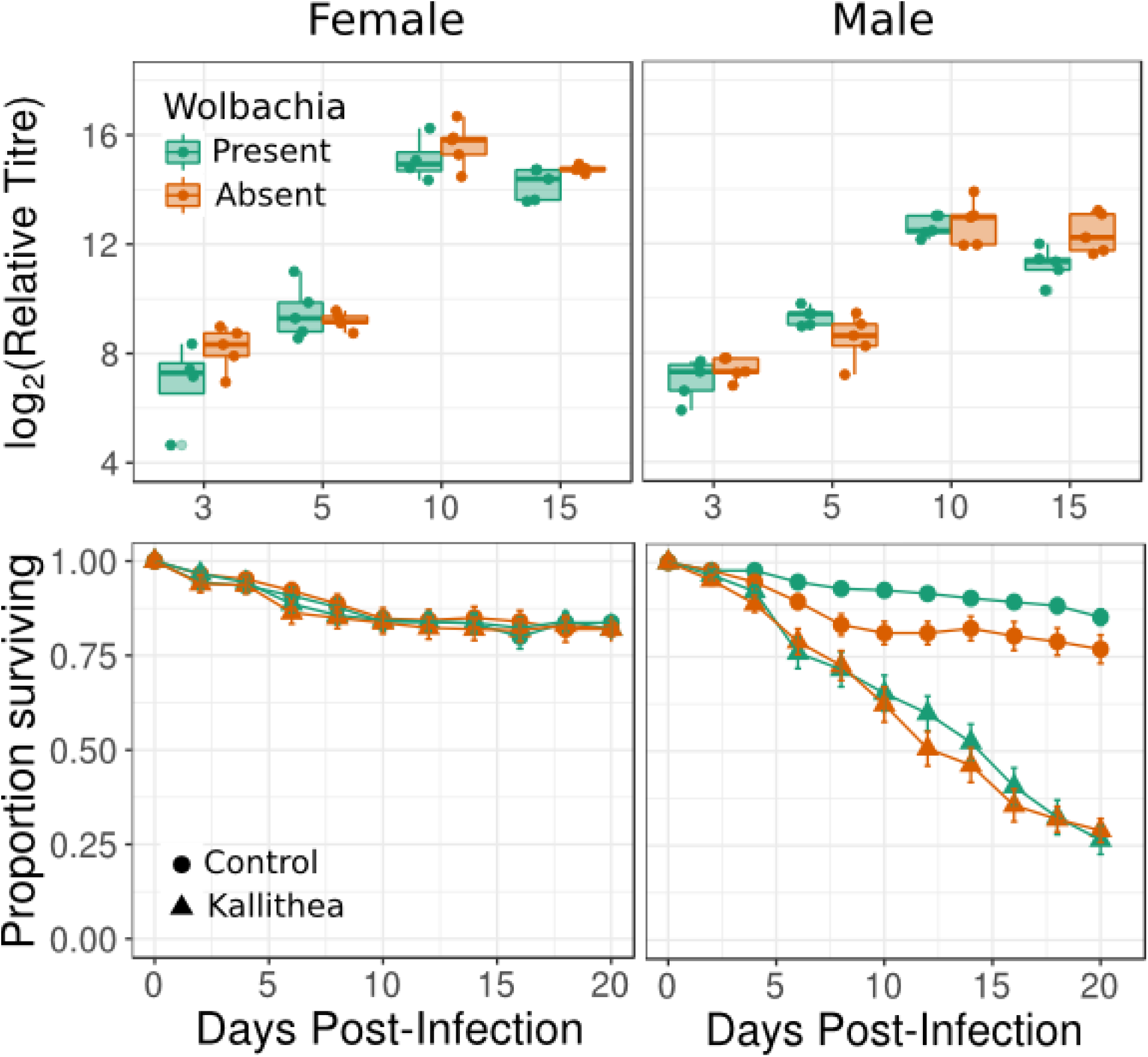
No effect of *Wolbachia* on KV growth or KV-induced mortality. Upper panels: Log-transformed relative viral titre in *Wolbachia* positive (green) or negative (orange) *OreR* female and male flies. Lower panels: mortality curves for gradient control-injected (circle) or KV-injected (triangle) *OreR* female and male flies, with or without Wolbachia.

**Figure S2:**
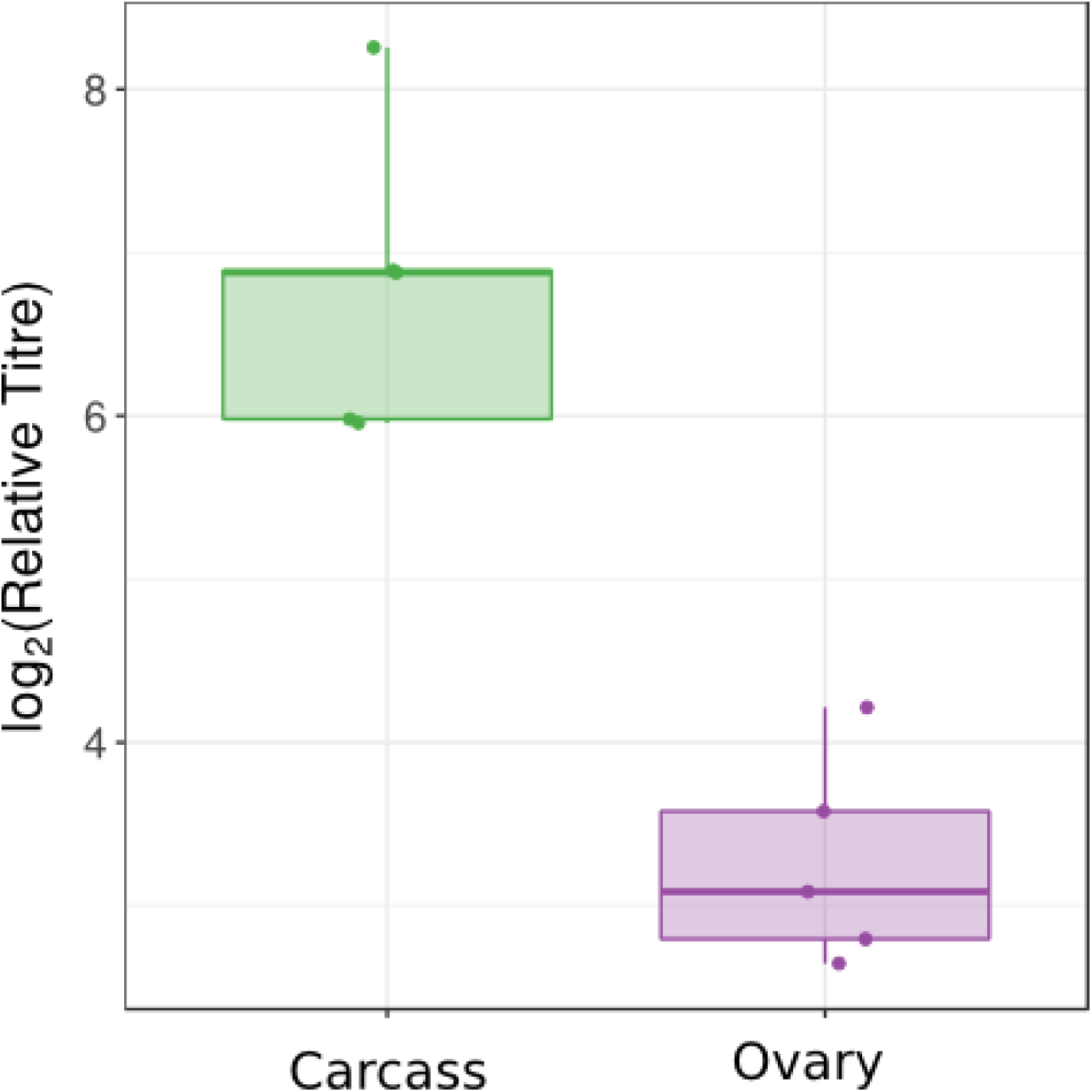
KV enriched in carcass relative to ovary. Females had higher viral titres in non-ovary tissues at 3 DPI. However, this could be affected by the route of infection, and the average ploidy of each sample.

**Figure S3:**
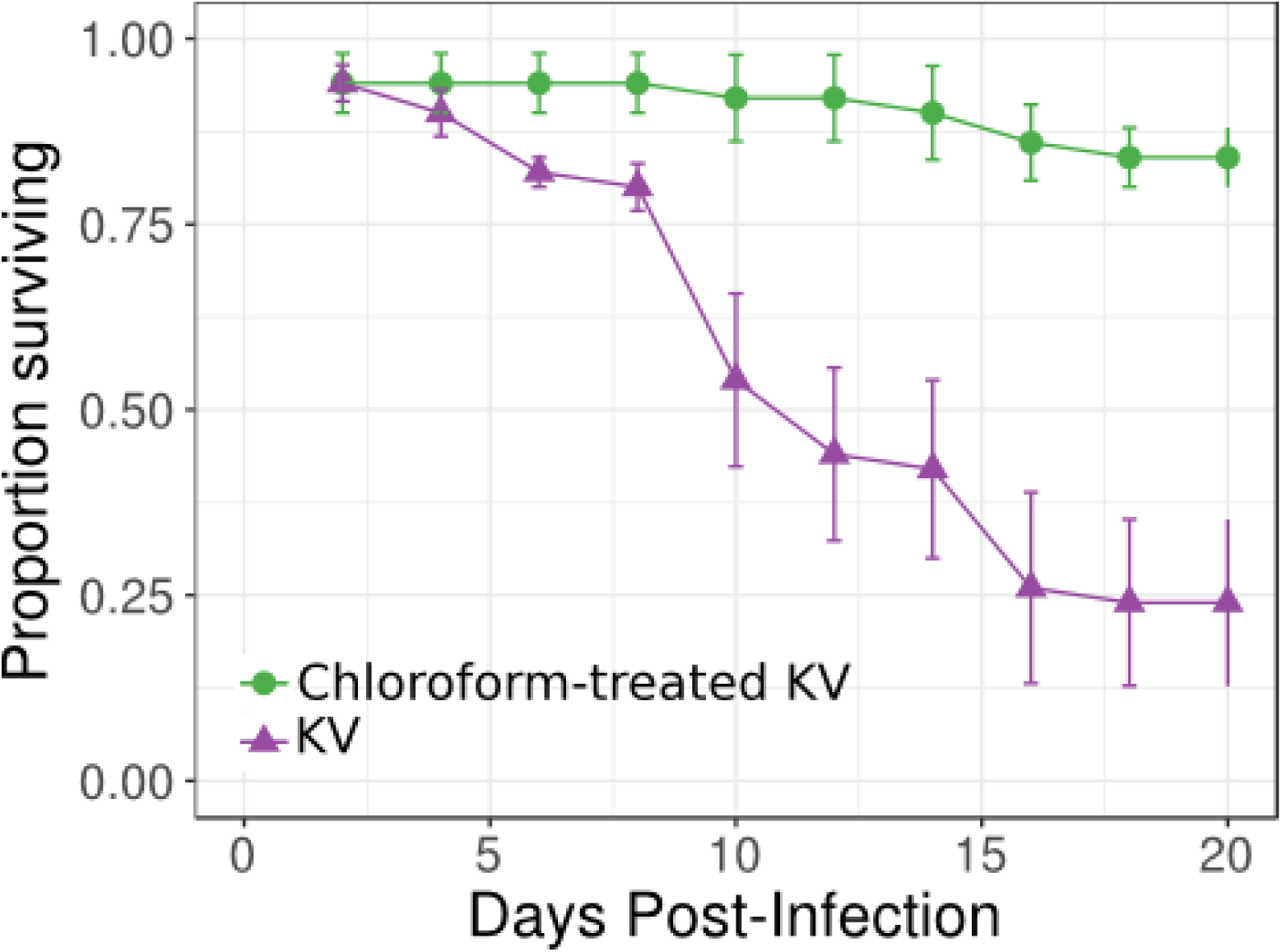
KV-induced male mortality is not caused by contaminating RNA viruses. Chloroform-treatment is expected to inactivate enveloped viruses such as KV, but unenveloped viruses (including most +ssRNA viruses) are expected to retain infectivity. We confirmed mortality following KV infection was not caused by contaminating DAV by comparing injection of the KV isolate with (green) or without (purple) inactivating chloroform treatment.

**Figure S4:**
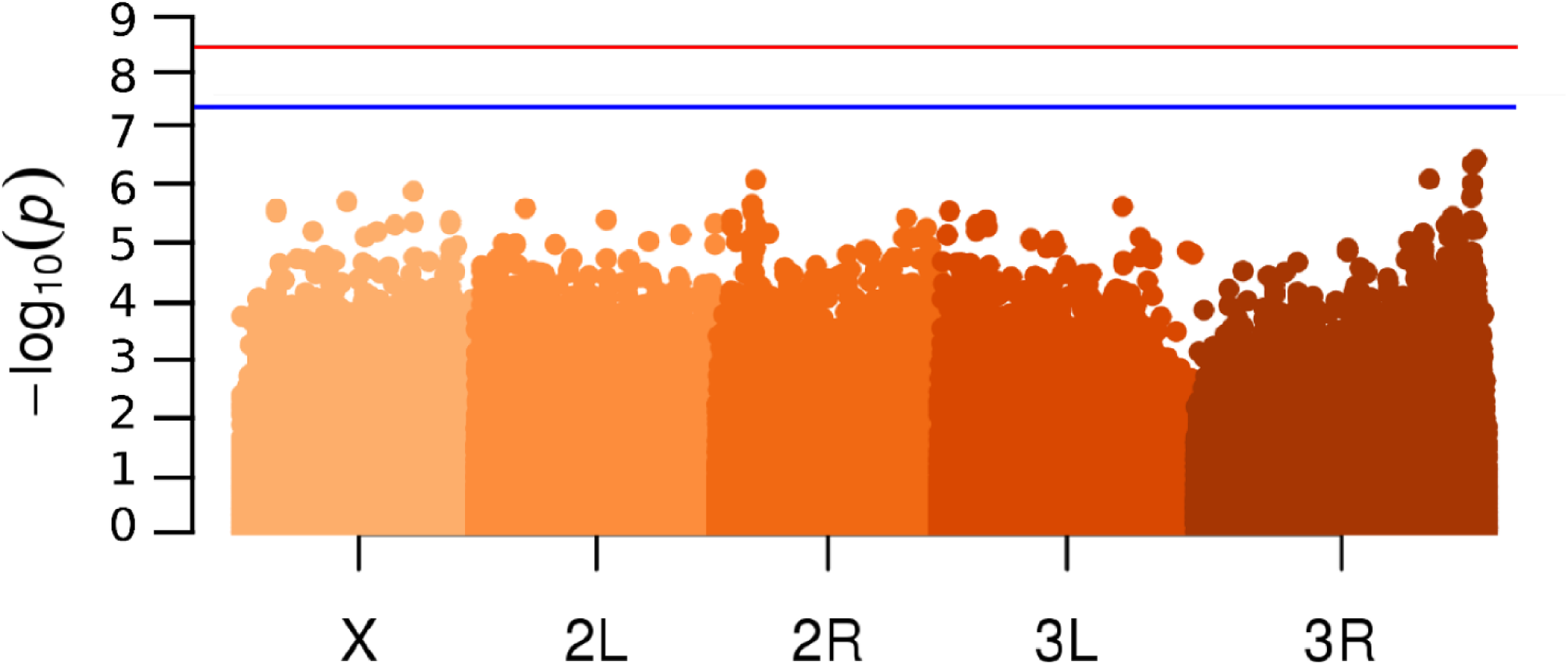
Sex-by-genotype interaction significance manhattan plot. No polymorphism had a significant effect on sex-specific mortality. The blue line denotes p_ran_ = 0.05 and the red line is p_ran_ = 0.01.

**Figure S5:**
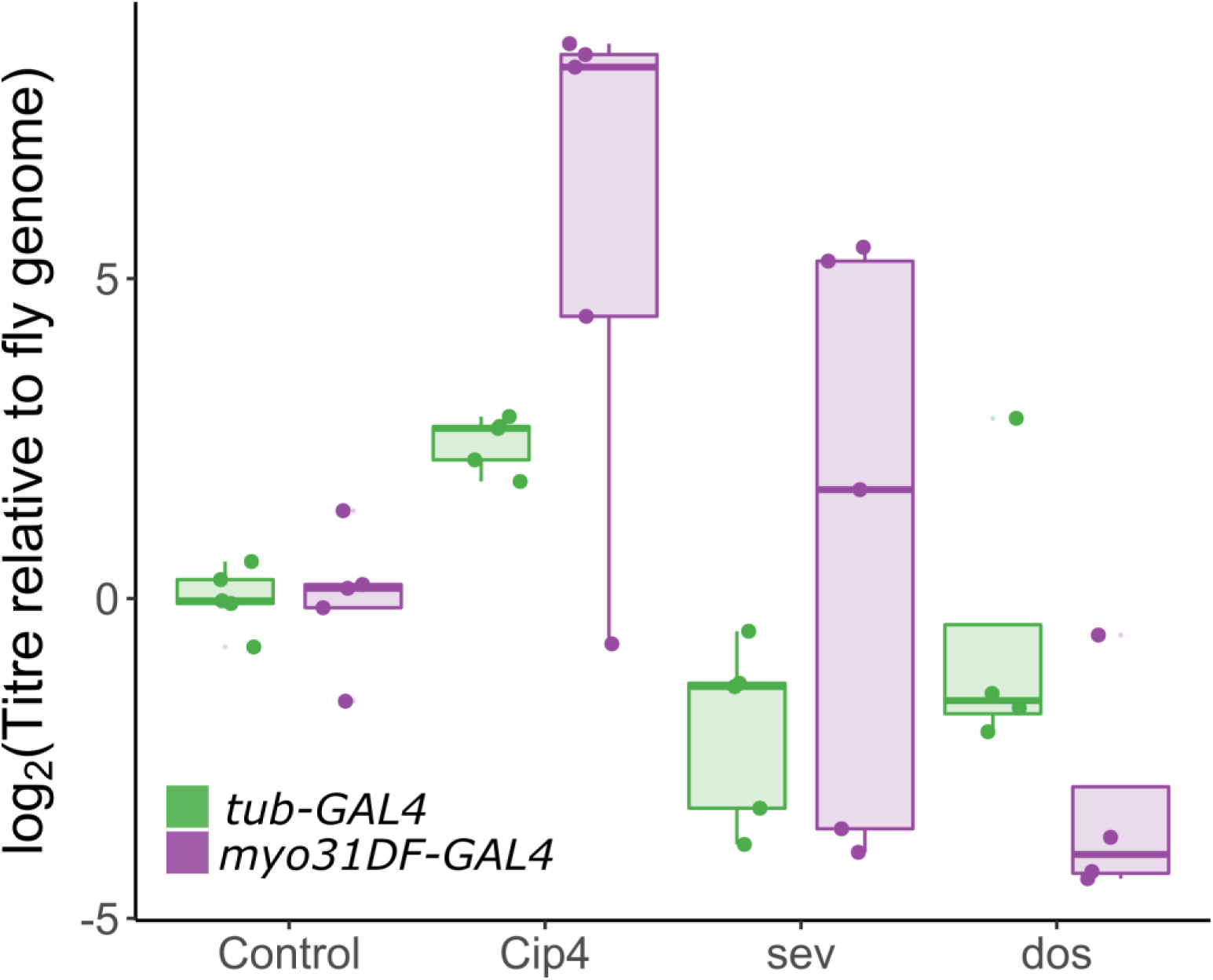
Virus titre in other confirmed GWAS knock-down lines. KV titre was measured in flies expressing a foldback hairpin targeting 18 genes identified in the GWAS, using GAL4 lines that knock each down in either the whole fly (*tub-GAL4*, green) or specifically in the gut (*myo31DF-GAL4*, purple). Only those causing a significant increase in titre relative to other knock-down lines (e.g. Figure 5) are shown here. Note that the cross between *myo31DF-GAL4* and *CG12821^IR^* was inexplicably lethal, and that titre was highly variable in some of the other *myo31DF-GAL4* crosses.

**Figure S6:**
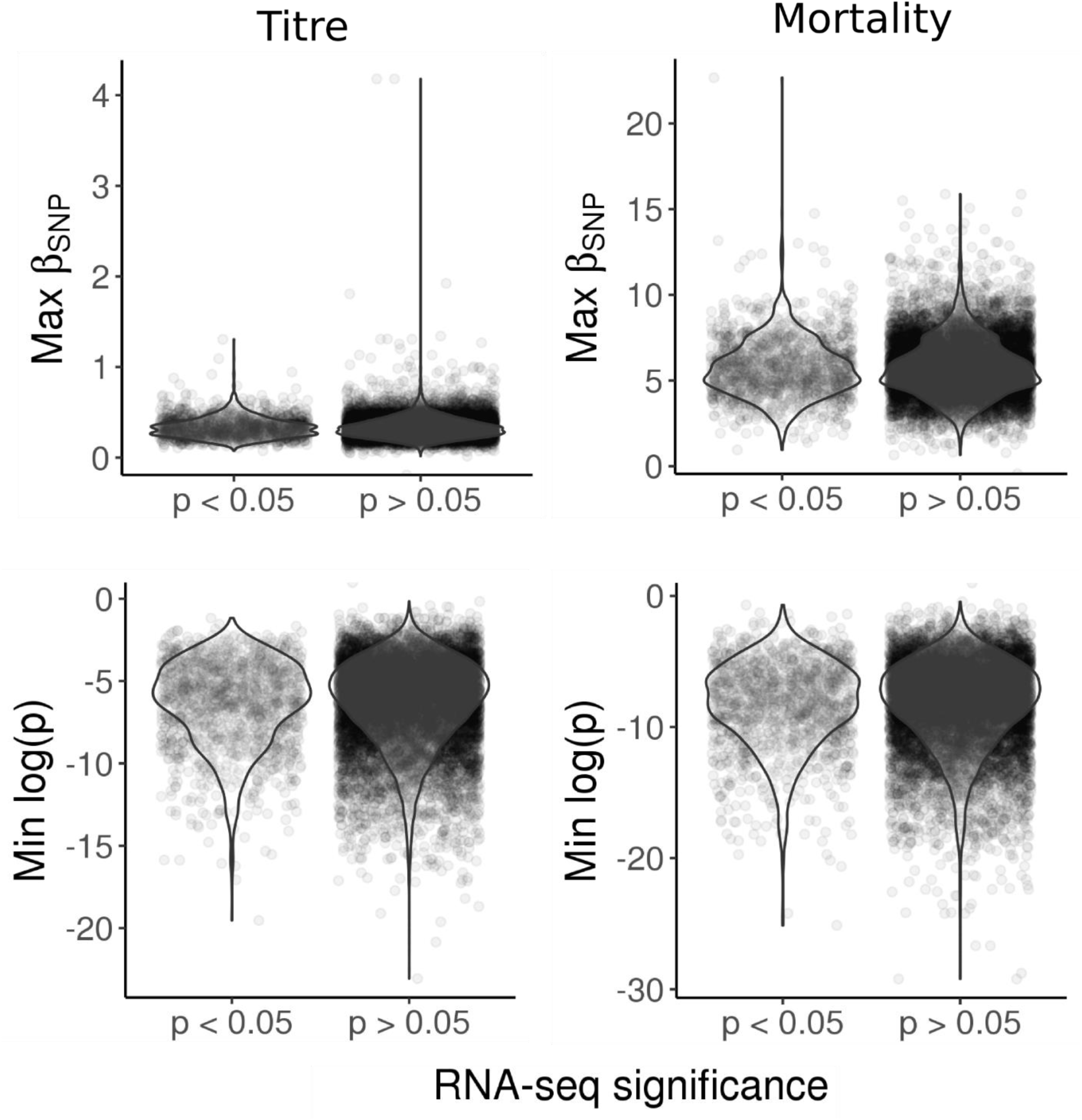
No enrichment of KV infection-associated SNPs in KV-responsive genes. Genes were split into KV-responsive and KV-unresponsive genes based on the RNA sequencing differential expression analysis (p < 0.05). The largest effect size (max β^SNP^) and lowest p-value was recorded for each gene in each GWAS, and compared between the KV-responsive and unresponsive genes. We find no significant difference between any comparisons.

**Figure S7:**
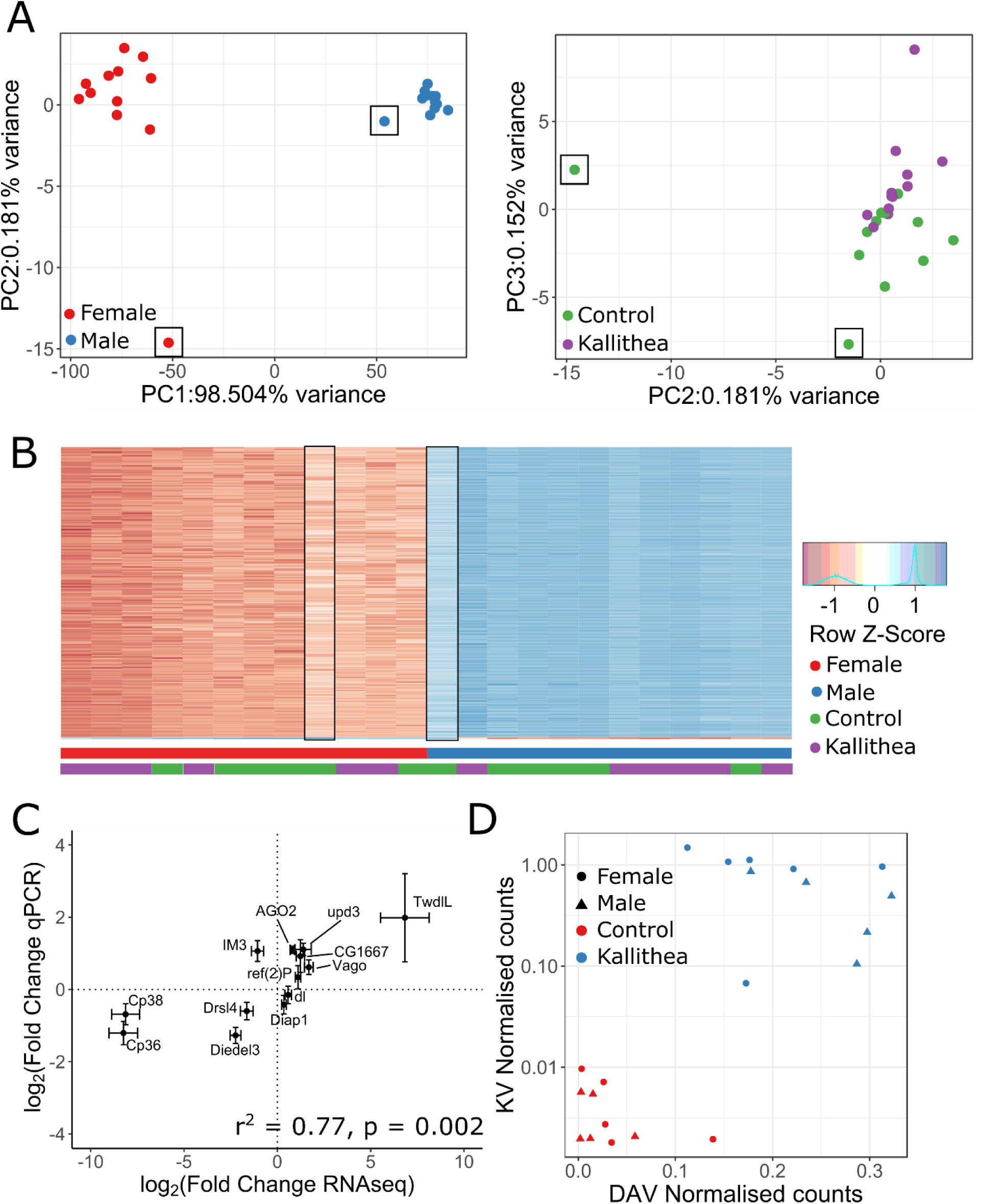
RNA sequencing: Library quality and verification. (A) The first three principal components of read counts per gene in RNA-sequencing data, plotted such that each library is represented by a point. Males (blue) and females (red) are separated on PC1. Control-injected (green) and KV-injected (purple) are separated on PC3. For (B), we clustered libraries based on expression of the 1000 most variable genes, where each row on the heatmap is a gene, and the columns are libraries. Together, these analyses identified two possible outlier libraries, which were excluded (black rectangles in A and B). (C) We selected 13 well-studied immune genes and genes with a clear phenotype association (e.g. chorion proteins), distributed across the range of differential expression values, for qPCR verification. Using 5 independent biological replicates from the outbred DGRP population, we confirmed that differential expression for these genes was highly correlated between qPCR and RNA-seq (r^2^ = 0.77, p = 0.002). (D) We found low-level DAV contamination in our RNA-sequencing experiment. Shown is the relationship between DAV viral titre and average KV gene expression, where each point is the number of reads mapping to KV and DAV for each library, normalised by library size factor and genome length. KV is plotted on a log_2_ scale.

**Figure S8:**
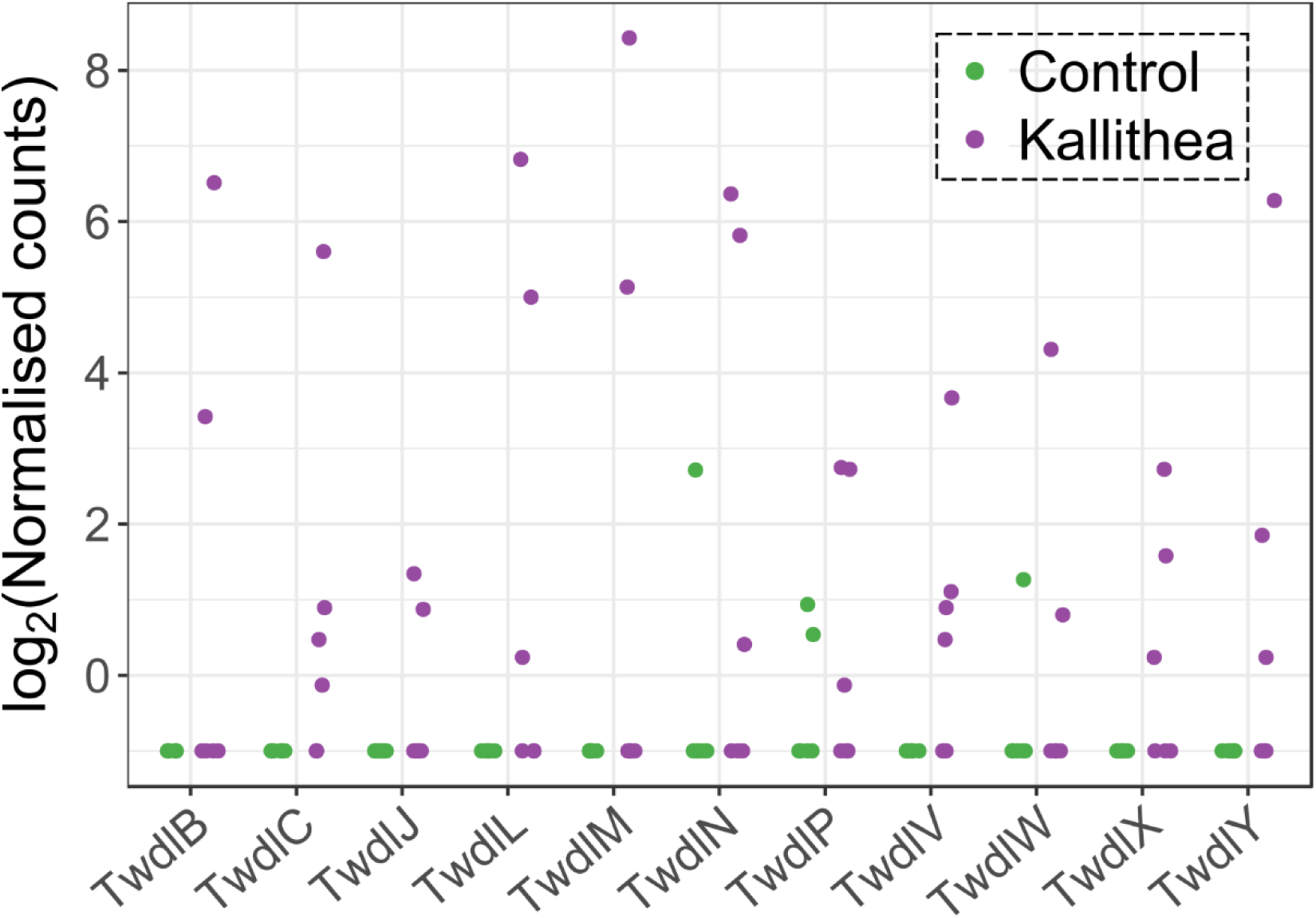
Variable differential expression of the Tweedle gene family. A subset of KV-infected (purple) vials showed very high expression of Tweedle genes, whereas these were mostly unexpressed in control (green) adult flies.

**Figure S9:**
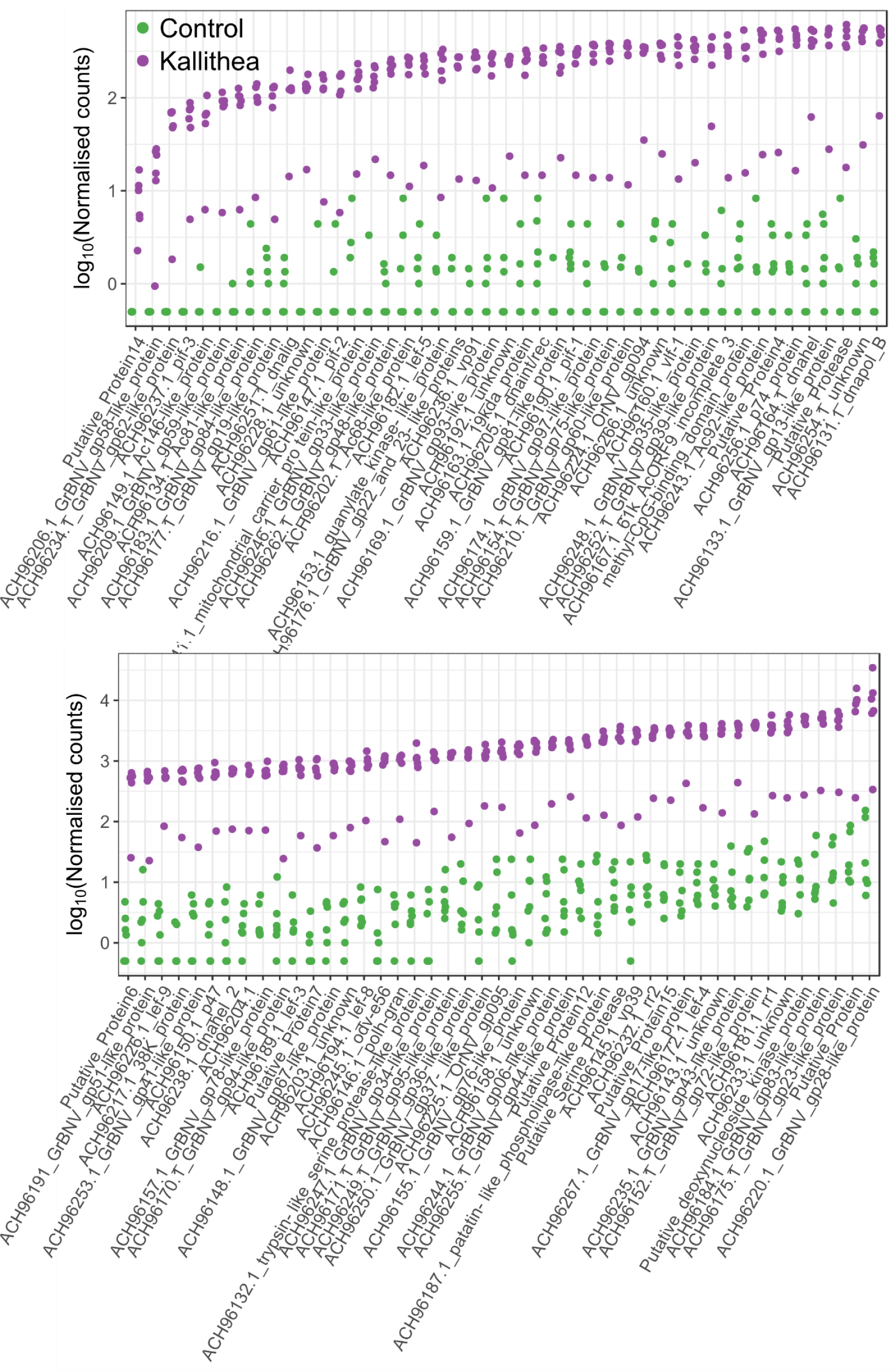
Expression of Kallithea Virus genes. Most KV genes are expressed at 3 DPI. One KV-injected vial of flies had a lower level of infection. Control libraries also showed mapping to KV genes, most likely due to a low level (<0.5%) of barcode switching among libraries run together. The lower panel is a continuation of the upper panel.

Table S1: Significant GWAS hits

Table S2: Significant GO term enrichment for GWAS hits

Table S3 Differentially expressed genes

Table S4 Significant GO terms for DE genes

